# Parameter inference on brain network models with unknown node dynamics and spatial heterogeneity

**DOI:** 10.1101/2021.09.01.458521

**Authors:** Viktor Sip, Spase Petkoski, Meysam Hashemi, Timo Dickscheid, Katrin Amunts, Viktor Jirsa

## Abstract

Model-based data analysis of whole-brain dynamics links the observed data to model parameters in a network of neural masses. In recent years a special focus was placed on the role of regional variance of model parameters for the emergent activity. Such analyses however depend on the properties of the employed neural mass model, which is often obtained through a series of major simplifications or analogies. Here we propose a data-driven approach where the neural mass model needs not to be specified. Building on the recent progresses in identification of dynamical systems with neural networks, we propose a method to infer from the functional data both the neural mass model representing the regional dynamics as well as the region- and subject-specific parameters, while respecting the known network structure. We demonstrate on two synthetic data sets that our method is able to recover the original model parameters, and that the trained generative model produces dynamics resembling the training data both on the regional level and on the whole-brain level. We further apply the method to resting-state fMRI data from Human Connectome Project. We find that to achieve best fit, the model needs two dimensional state space and three regional parameters, one of which is strongly correlated with the map of genetic expression in human brain. The present approach opens a novel way to the analysis of resting-state fMRI with possible applications in understanding the changes of whole-brain dynamics during aging or in neurodegenerative diseases.

## 1. Introduction

One avenue for analysis of resting-state functional magnetic resonance imaging (fMRI) is the use of computational models of large-scale brain network dynamics (1, 2). A general goal of this approach is to relate the observed brain activity to the dynamical repertoire of the computational model, possibly via identification of optimal model parameters, leading to a better mechanistic interpretation of the observations. Such models are network-based, where the nodes represent brain regions and the edges the structural connections between them. Models constrained by individual brain imaging data are referred to as virtual brains and typically use diffusion-weighted imaging data for the edges and neural mass models for the local dynamics of a brain region. Neural masses are low-dimensonal models of neuronal population activity.

When linking the models with the observations, until recently studies focused only on a small number of parameters - such as the global coupling strength - due to the computational costs associated with the exploration of a high-dimensional parameter space. In recent years, however, several works utilized the whole-brain modeling framework in order to explore the role of spatial heterogeneity of model parameters. Specifically, the studies found that the whole-brain models can better reproduce the features of resting-state fMRI when the regional variability is constrained by the MRI-derived estimates of intracortical myelin content (3), functional gradient (4), or gene expression profiles (5), and similar regional variability was found even without prior restrictions (6).

Neural mass models employed in these studies (such as the dynamic mean field model of conductance-based spiking neural network (7) or Hopf bifurcation model of neural excitability (8)) are derived through a series of major simplifications or built upon loose mathematical analogies. It can thus be questioned to what degree the dynamical structure embodied in these models is sufficient to capture the essential elements of the neural dynamics manifesting in the observed data. Would two different neural mass models lead to the same conclusions, or do the results strongly depend on the exact model form? Such questions are not yet sufficiently answered.

Meanwhile, novel techniques to learn the models of nonlinear dynamical systems from the data itself are being developed and applied in various fields of physical and life sciences (9–13), including in neuroscience on all scales (14–16). The common assumption in these approaches is that the observed data are generated by an unknown dynamical system of reasonably low dimensionality, which can be represented with a flexible artificial neural network. The parameters of this network are learned during training, so that it best reproduces the data.

These developments raise the question whether a similar approach can be applied in the context of whole-brain modeling: Can we learn a dynamical system representing a neural mass at each node of a large-scale brain network? Such approach would allow to side-step the issue of reliance on a specific neural mass models which lie at the heart of the large-scale modeling, and instead extract this model directly from the functional data. That is what we aim to investigate in this work. Using the known network structure, derived from diffusion-weighted imaging, and the observed resting-state fMRI, we infer the dynamical system representing the neural masses in the nodes of the network. To account for the regional and subject heterogeneity, we allow this (initially unknown) neural mass model to depend on a region-specific and subject-specific parameters. These parameters we also infer from the observations together with the model, obtaining the map of dynamically-relevant parameters (Fig. 1).

**Figure 1:**
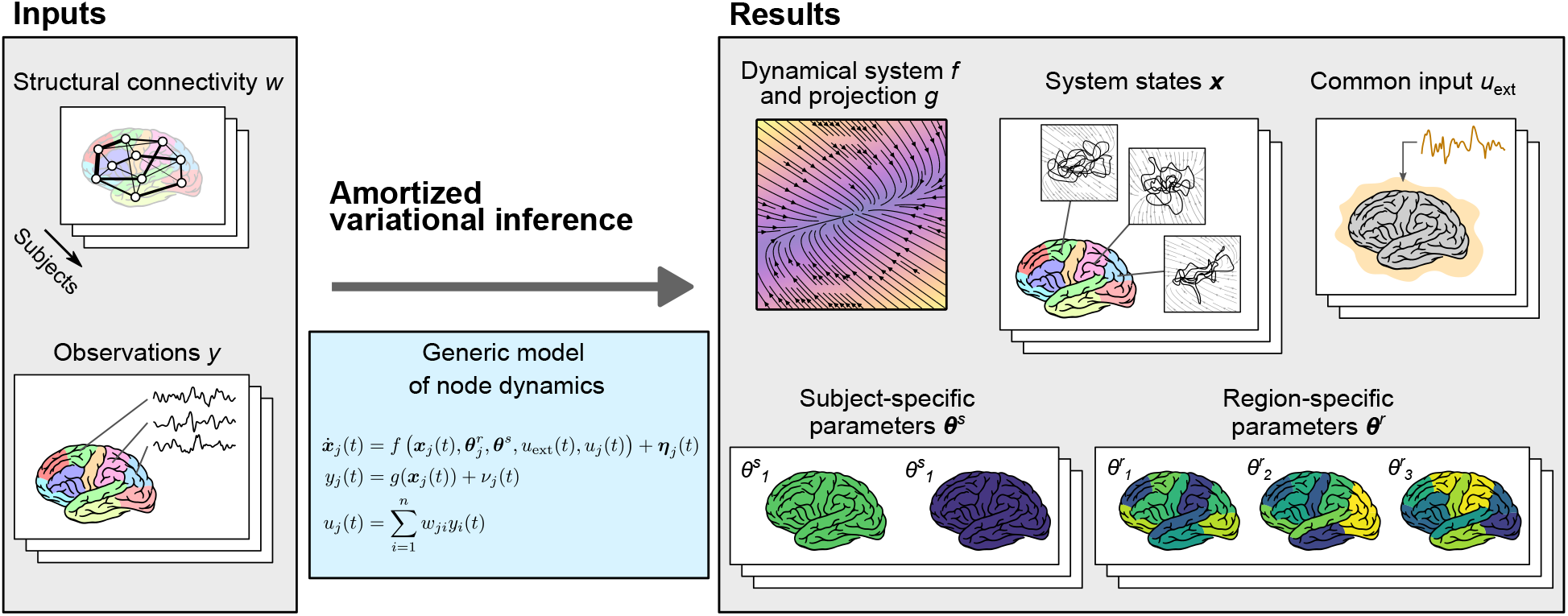
Conceptual overview of the method. The method allows to perform a parameter inference for network models of brain dynamics, where the dynamical model of every node (or brain region) is initially unknown. As input (left) it expects structural connectivity matrices *w* for cohort of multiple subjects, and corresponding observations of brain activity ***y*** (such as parcellated resting-state fMRI). Constrained by the structure of a generic model of regional dynamics (middle), it learns the dynamical model of node dynamics *f* and the state-to-observation projection model *g*. The dynamical model *f* is shared for all subjects and regions, but it depends on subject-specific parameters ***θ***^*s*^ and region-specific parameters ***θ***^*r*^. These too are inferred from the data, together with the hidden states of the system ***x***, and the subject-specific external input *u*_ext_, shared by all regions in a given subject. All of the system states, subject- and region-specific parameters, and the external input are inferred probabilistically as normal distributions, that is, we infer the mean and variance for each parameter.

To do so, we utilize the framework of amortized variational inference, or variational autoencoders (17), inspired in particular by its application for inferring neural population dynamics (14) and for dynamical systems with hierarchical structure (11). In brief, our system is composed of an encoding network, mapping the observed time series to the subject- and region-specific parameters and to the trajectory in the source space, a neural network representing the dynamical system, and the observation model acting as the decoder from the source to the observation space. These are jointly trained to maximize the evidence lower bound, so that the predictions of the trained model closely resemble the original data.

In this work we test our method on two synthetic data sets, generated with the two models commonly used in large-scale brain modeling: the mean field model of conductance-based spiking neural network, or mean field model for short (7), and the Hopf bifurcation model (8). For both test cases we use a cohort of eight subjects with realistic structural connectomes, and with model parameters varying across subjects and brain regions. We show that the trained generative model can reproduce many features of the original data set, and we demonstrate that the method can extract regional and subject-specific parameters strongly related to the original parameters used for the simulation. We further apply the method to resting-state fMRI data from Human Connectome Project (18). We find that to achieve best fit, the model needs two dimensional state space and three regional parameters, one of which is strongly correlated with the map of genetic expression in human brain.

## 2. Results

The results section consists of three parts. In the first part, we introduce the general ideas of the developed method; its detailed description can be found in Methods. In the second part, we validate the proposed method on synthetic data, that is, on data generated by computational models with known parameters. In the third part we apply the method on resting-state fMRI data from human subjects and analyze the results.

### 2.1. Amortized variational inference for networks of nonlinear dynamical systems

We follow the general framework of large-scale brain network modeling, and we assume that for a specific subject the observations *y*_*j*_(*t*) of a brain region *j* are generated by a dynamical system

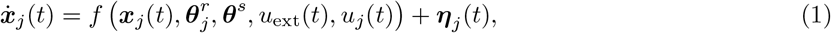

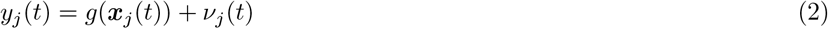

where 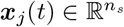 is the state at time 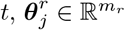 and 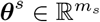 are the region-specific and subject-specific parameters.

The term *u*_ext_(*t*) is the external input, shared by all regions of a single subject, and

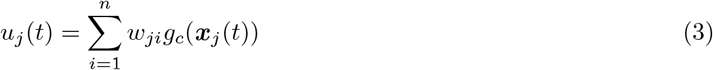

is the network input to region *j* with 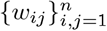 being the structural connectome matrix of the network with *n* nodes. The functions *f, g*, and *g*_*c*_ are initially unknown, and ***η***_*j*_ (*t*) and *ν*_*j*_(*t*) is the system and observation noise, respectively (Fig. S1A).

From the observed time series of multiple subjects we wish to infer both the evolution function *f* and observation function *g* (and coupling function *g*_*c*_, which for simplicity we assume is identical to *g*), shared across the subjects, as well as region- and subject-specific parameters 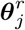 and ***θ***^*s*^ and the time-dependent external input *u*_ext_. To do so, we adopt the general framework of amortized variational inference (17) with hierarchical structure in parameters (11) (Fig. S1B). We consider the states ***x***_*j*_, the parameters 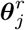 and ***θ***^*s*^, and the external input *u*_ext_ the latent variables, and we seek their approximate posterior distribution represented by multivariate Gaussian distributions. In the spirit of amortized variational inference, we do not optimize their parameters directly, but through encoder functions *h*_1_, *h*_2_, *h*_3_, and *h*_4_ which transform the data to the latent variables (system states, regional and subject parameters, and external input respectively).

The assumption that the observation and coupling functions are identical, *g* ≡ *g*_*c*_, allows us to effectively decouple the network problem to uncoupled regions with known network input, and so we can consider timeseries of one region of one subject as a single data point. We represent the nonlinear function *f* with a generic artificial neural network, and function *g* as a linear transformation. The inference problem is ultimately transformed into optimization of the cost function, evidence lower bound (ELBO), which is to be maximized over the weights of *f, g, h*_1_, *h*_2_, *h*_3_, and *h*_4_, and over the variances of the system and observation noise. After the optimization, we obtain the description of the dynamical system in terms of functions *f* and *g*, probabilistic representation of the regional and subject parameters 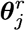 and ***θ***^*s*^ and of the external input *u*_ext_, as well as projections of the observations in the state space ***x***_*j*_. The inferred parameters 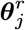 and ***θ***^*s*^ will not have a mechanistic meaning; however, they can provide a measure of (dis)similarity of the regions and subject, and they can be interpreted via the inferred dynamical system *f*.

### 2.2. Validation on synthetic data

#### Evaluation workflow

We test the proposed method on two synthetic data sets, where the data are generated by models commonly used in whole-brain modeling. First is the Hopf bifurcation model (8), shown on Fig. 2. That is a two-equation neural mass model, where depending on the value of the bifurcation parameter *a*_*i*_ the dynamics is either noise-driven around a stable fixed point (for *a*_*i*_ *<* 0) or oscillatory with frequency *f*_*i*_ (for *a*_*i*_ *>* 0). In the synthetic data set, these two parameters are randomly varied across regions. Second model is the parametric mean field model (pMFM 7), shown on Fig. 3. That is an one-equation model, and depending on the network input, it can be pushed into monostable down- or up-state, or a bistable regime. The switching between the states is noise driven, and we vary the noise strength across brain regions.

**Figure 2:**
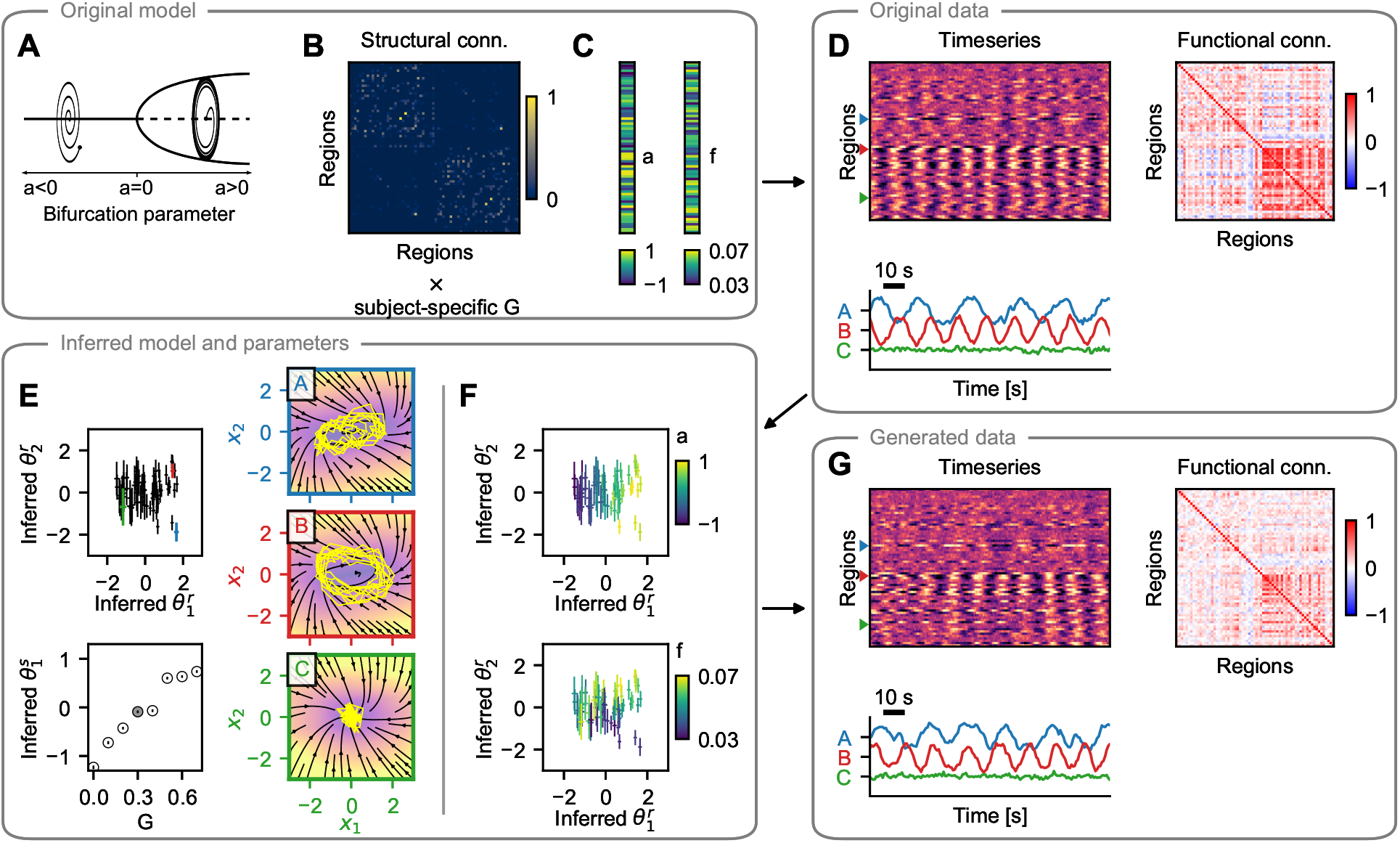
Hopf model test case: example subject. (A-C) The training data are simulated using a network model of brain dynamics, where in each node a Hopf neural mass model is placed (A). The nodes are coupled through a connectome derived from diffusion-weighted imaging (B) scaled by a subject-specific coupling parameter *G*. The values of bifurcation parameter *a*_*i*_ and intrinsic frequency *f*_*i*_ vary across brain regions (C). (D) Timeseries generated with the original model with three examples (bottom) and the calculated functional connectivity (right). (E) Inferred regional parameters for all regions (top left, example nodes highlighted in color) and inferred subject-specific parameter (bottom left, in gray among parameters for all subjects in the dataset). The span of the crosses/lines corresponds to two standard deviations of the inferred Gaussian distribution. In the bottom panel circles are added for visual aid due to the small standard deviations. The inferred dynamics in state space of the three example nodes are on the right. The vector field is evaluated assuming zero network input and using the inferred region- and subject-specific parameters. Background color represents the velocity magnitude, yellow lines are exemplary simulated time-series of the node activity when embedded in a network. (F) Inferred regional parameters colored by the ground truth values of the bifurcation parameter *a*_*i*_ (top) and frequency *f*_*i*_ (bottom). The bifurcation parameter correlates with inferred 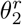, while frequency correlates with 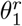, but only for regions in the oscillatory regime, i.e. where *a*_*i*_ *>* 0. (G) Timeseries generated with the trained model and using the inferred parameters. Important features of the data are preserved both the level of single regions (amplitude, frequency) as well as on the network level (functional connectivity).

**Figure 3:**
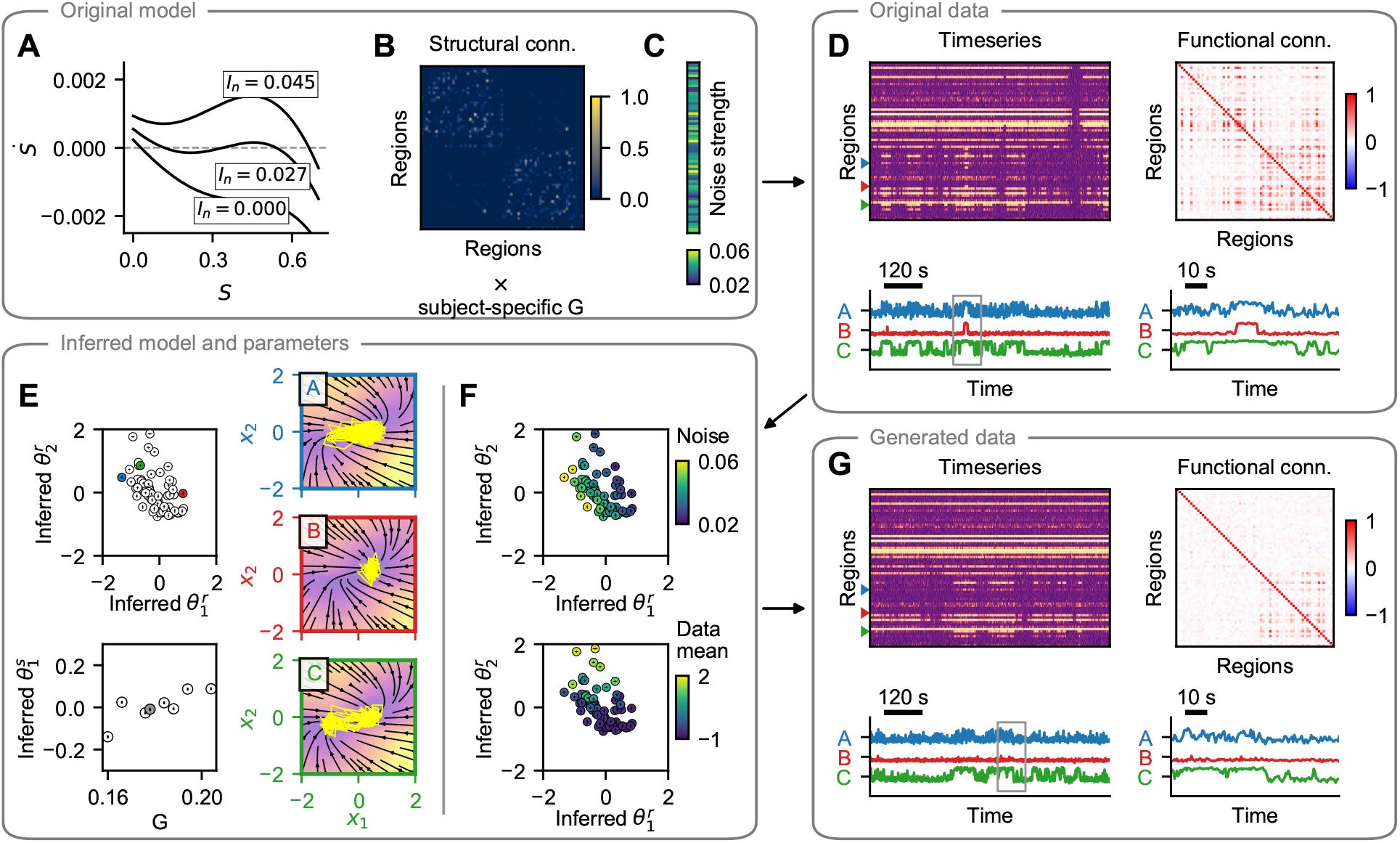
Parametric mean field model test case: example subject. Layout same as in Fig. 2. (A-C) The training data are simulated using a network of pMFM neural masses. Depending on the network input, these can be forced into the monostable regime (down or up state) or into the bistable regime. The dynamics is noise driven, with noise strength varying across regions. (D) Timeseries generated with the original model and the functional connectivity. Three examples shown in the bottom panel, with a window of hundred seconds on the right. (E) Inferred regional parameters (top left) and subject-specific parameter. Circles are added for visual aid due to the small standard deviations. The inferred dynamics in state space of the three example nodes are on the right. The vector field is evaluated assuming zero network input and using the inferred region- and subject-specific parameters. Background color represents the velocity magnitude, yellow lines are exemplary simulated time-series of the node activity when embedded in a network. (F) Inferred regional parameters colored by the ground truth values of the noise strength parameter (top); the original parameter is encoded along the diagonal of the inferred parameters. Bottom panel shows coloring according to the mean of the original timeseries, which does not represent an original model parameter, rather a data feature. (G) Timeseries generated with the trained model and using the inferred parameters. Region-specific features (switching between states, noisiness) are well preserved. Structure of the regional correlations is also reproduced, but the correlations are weaker compared to the original.

Both models are used to generate synthetic data for eight subjects, each with individual structural connectome containing 68 cortical regions of the Desikan-Killiany parcellation (19). The connectome is scaled by the global coupling strength *G* which we set to increase linearly across subjects for the Hopf model, or which we set to the optimal value (in terms of highest produced functional connectivity), different for every subject, with pMFM.

To establish the performance of the described method, we proceed as follows. First, we simulate the data with the original model and random values of regional parameters (Fig. 2D and Fig. 3D). Next, using the whole data set of eight subjects, we train the model, obtaining at the same time the trained generative model described by the function *f* of the dynamical system, and also the probabilistic representation of subject- and region-specific parameters (Fig. 2E and Fig. 3E). Taking random samples from the posterior distributions of the parameters, and using random system and observation noise, we repeatedly generate new timeseries using the trained model (Fig. 2G and Fig. 3G).

We evaluate the quality of the trained model based on the following criteria. First, we establish whether the inferred parameters are related to the original parameters of the model (Figs. 2F, 3F, 4A,E). Second, we wish to evaluate whether the features of the generated timeseries resemble those of the original timeseries, both on the regional level (Fig. 4B,F) and on the network level (Fig. 4C,D,G,H). We explore these aspects in the following paragraphs.

**Figure 4:**
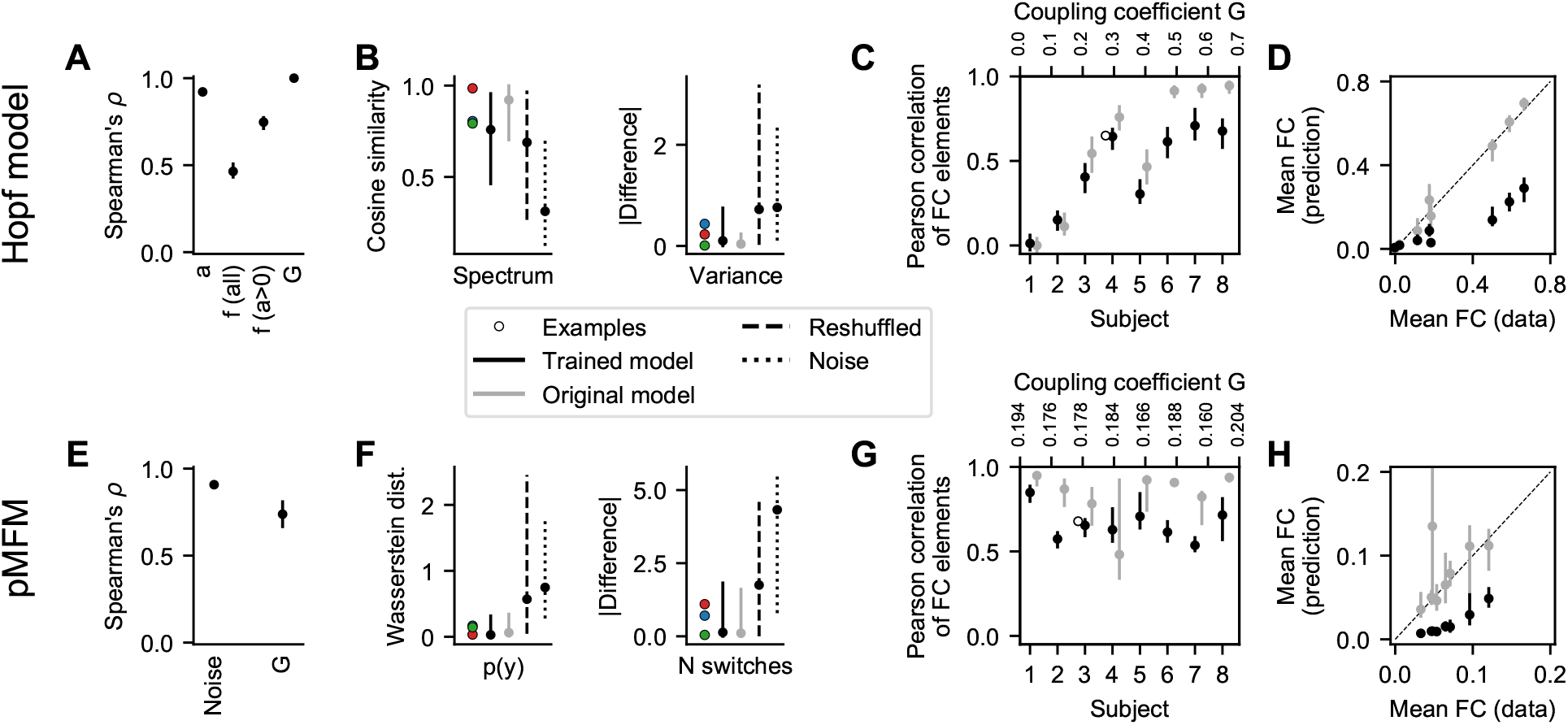
Quantitative evaluation of the synthetic test cases. Top row - Hopf model, bottom row - parametric mean field model. (A, E) Nonlinear correlation between the original parameters and the optimal projection of the inferred region-specific parameters (bifurcation parameter *a* and frequency *f* for Hopf model, noise strength for pMFM) and subject-specific parameter (coupling strength *G*). (B, F) Fit between the regional features of the original timeseries and those generated by the trained model. We show the cosine similarity of the timeseries spectra and the difference in variance for the Hopf model, and Wasserstein distance of the distributions in the observation space and the difference in logarithm of number of switches for pMFM. These are evaluated for the examples from Fig. 2 and Fig. 3, all timeseries generated by the trained model, and the surrogates described in the main text. (C, G) Fit between the functional connectivity of the original and generated timeseries. (D, H) Mean value of non-diagonal elements of functional connectivity matrices. For both models, the correlation strength is underestimated, even if the structure is preserved. In all panels, the bars show the (5, 95) percentile interval with the dot representing the median value. The statistics are computed from 50 samples of the posterior distribution for 8 subjects (grouped together in A, B, E, F) and 68 regions (for region-specific parameters and features). The statistics of the surrogate distributions using the original model are also calculated from 50 samples.

#### Inferred parameters encode the original model parameters

The example on Fig. 2 shows how are the original regional parameters encoded in the inferred parameters ***θ***^*r*^ for the Hopf model. The bifurcation parameter *a* is encoded in the inferred parameter 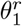 (upper panel), while the frequency *f* is encoded in 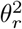 (lower panel). The latter is however true only for the regions in the oscillatory regime, i.e. with *a >* 0. That is not a deficiency of the proposed method: in the fixed point regime the activity is mainly noise-driven, and the value of the frequency parameter has small to negligible influence (see the example C on Fig. 2D). In other words, the parameter is not identifiable from the data. That is reflected in the inferred parameters. For the regions with *a >* 0 (or equivalently with 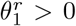) the inferred parameters 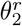 have low variance, and their mean encodes the original frequency parameter. For the regions with *a <* 0, however, inferred 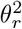 have high variance, close to the prior value 1, and overlapping distributions, indicating that not much information is encoded in 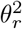 in this regime.

Also for the pMFM test case the noise strength parameter is well identified (Fig. 3F), however the second dimension of the region-specific parameter 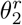 is used to encode the mean of the regional timeseries. Presumably, this is so that the parameter 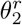 can compensate for the weaker network coupling, which we discuss later. For both examples the subject-specific coupling strength is encoded in the subject parameter ***θ***^*s*^ (Figs. 2E, 3E, lower panels).

The quantitative analysis of the goodness-of-fit is shown on Fig. 4A,E. To evaluate it, for each of the original parameters we first identified the direction in the parameter space along which the parameter is encoded by taking a linear regression of the posterior distribution means. Then, we repeatedly took samples from the posterior distributions of the parameters, projected them on the identified subspace, and calculated the nonlinear Spearman’s correlation coefficient *ρ*. For most parameters the values are close to the optimal value of 1, indicating that the original parameters are indeed accurately recovered in the inferred parameters. The exception is the frequency *f* due to the above discussed non-identifiability. If, however, we restrict the regions only to those where the bifurcation parameter is positive, the correlation markedly increases, as expected based on the discussed example.

On Fig. S2 we further evaluate how the goodness-of-fit changes with the increased coupling in the Hopf model. Presumably, as the coupling increases, the regional timeseries are more affected by the activity of the connected regions and less by its internal parameters, and it is thus more difficult to recover the original parameters from the data. Indeed, that is the trend that we observe both for the bifurcation parameter *a* and frequency *f* of the nodes in oscillatory regime.

#### Trained model reproduces the features of regional timeseries

A crucial test of the trained model is an evaluation whether the generated data resemble those used for the initial training. This resemblance should not be understood as reproducing the timeseries exactly, since they depend on a specific noise instantiation, rather that the features we consider meaningful should be preserved. For both test cases, we evaluate the similarity of two features. For the Hopf model with its oscillatory dynamics we evaluate the cosine similarity of the spectra of the original and generated timeseries, and the difference between the variance of the timeseries, since the variance differs greatly between the nodes in oscillatory and fixed-point regimes (Fig. 4B). For the pMFM, we compare the timeseries based on the distribution in the 1D observation space (that is, taking the samples collapsed across time) using the Wasserstein distance (called also Earth mover’s distance) of two distributions. Second feature of pMFM timeseries is the log-scaled number of switches between the up- and down-state, capturing the temporal aspect of the switching dynamics (Fig. 4F).

We evaluate the measures for 50 different noise instantiations, leading to 50 different time series for each region, obtaining a distribution of goodness-of-fit metrics. The same metrics are evaluated also for three surrogates: First is the original computational model, run with different noise instantiations. That provides an optimistic estimate of what can be achieved in terms of goodness-of-fit, considering that the features will necessarily depend on the specific noise instantiation used in the initial simulations. Second surrogate is obtained by randomly reshuffling the original data between regions and subjects. Third surrogate is simply white noise with zero mean and variance equal to one (which, due to the data normalization, is equal to the mean and variance of the original data set taken across all subjects and regions).

In most measures, the trained model performs comparably or slightly worse than the original model and markedly better than the surrogates. The exception is cosine similarity of the spectra with the Hopf model. That is due to the very strong coupling in some subjects (for coupling coefficients *G* ≥ 0.5) leading to close to homogeneous activity across brain regions, and thus comparable performance of the reshuffling surrogate.

#### Functional network structure is reproduced, but with lowered strength

Just as the well trained model should be able to reproduce the features of the original data on the level of single regions, it should also be able to reproduce the relevant features on the network level. Specifically, we evaluate how well is the functional connectivity reproduced. In general, functional connectivity (FC) quantifies the statistical dependencies between the time series from brain regions. While there are multiple ways to measure it, the most ubiquitous is the linear (Pearson’s) correlation of the time series, which we use here as well. This static FC captures the spatial structure of statistical similarities, however, it has its limitations, notably it ignores the temporal changes in FC structure (20, 21).

The examples for both investigated models indicate that the FC structure is indeed well reproduced, but with lower strength, particular in the case of pMFM example (Figs. 2G and 3G).

This is further analyzed for all subjects on Fig. 4, and visualized on Fig. S3 and Fig. S4. For the Hopf model, the coupling coefficient was increased between subjects. For low coupling values, the FC structure is not reproduced (as measured by Pearson correlation between the non-diagonal elements of original FC and trained model FC; Fig. 4C). That is however true also for the original model due to the FC elements being close to zero and noise-dependent. For stronger coupling, the structure is preserved better, although the trained model plateaus around values of 0.7 for the correlation between the FC matrices, even when the correlations between the original model increases further. The comparison of the mean value of non-diagonal FC elements furthermore reveals that the strength of the correlations is considerably underestimated with the trained model (Fig. 4D).

For the pMFM, the coupling coefficient was set to optimal value (in the sense of maximal FC), specific to each subject. Also there we can see well reproduced structure of the correlations (Fig. 4G), although too with reduced strength (Fig. 4H).

These results indicate that while the trained model can discover the existence of the network coupling, it systematically underestimates its strength. Given that in the pMFM the strength of the network input can shift a single neural mass from the monostable down-state to bistable regime and to monostable up-state, the underestimated coupling leads to the necessity of utilizing the regional parameter to compensate for the missing coupling (Fig. 3F).

#### Large perturbations of the connectome lead to reduced performance

To assess the influence of the inexact structural connectome on the goodness-of-fit, we have trained the model on the pMFM data set with perturbed connectomes. That is, instead of the original connectivity matrix *W* we have trained the model with *W*_*ϵ*_ = *W* + *ϵA*, where *A* is matrix with elements drawn from standard normal distribution, and *ϵ >* 0 is the perturbation magnitude. In addition we have also used a log-scaled connectivity matrix.

Fig. S5 shows how are the indicators of goodness-of-fit from Fig. 4 modified by these perturbed connectomes. High perturbation magnitudes reduces the recovery of regional and subject parameters (Fig. S5A,B) as well as the similarity of the generated functional connectivity (Fig. S5E). The regional features, on the other hand, are reproduced similarly well even for large perturbations (Fig. S5C,D). Using the log-scaled connectome has a similar negative effect, although less pronounced.

### 2.3. Application on human resting-state fMRI

We applied the developed method to human resting-state fMRI data obtained from Human Connectome Project (18). For eight subjects we have analyzed fMRI time series from one session (1200 seconds), processed with HCP pipeline and further denoised by DiCER method (22), parcellated into 68 cortical regions of Desikan-Killiany parcellation (19). Where not mentioned otherwise, the results presented are obtained with state space dimension *n*_*s*_ = 3, regional parameter dimension *m*_*r*_ = 3, subject parameter dimension *m*_*s*_ = 2, and for the model variant with external input and with standard preprocessing of the structural connectomes (see Methods for details).

#### External input improves the FC fit

We started the analysis by evaluating the choice of the model, in particular, what effect has the presence of an external input in the model, and how does preprocessing of the structural connectome affect the results. Fig. 5A-E shows an example of single subject data processed by the models without and with an external input, while Fig. 5F shows the group results.

**Figure 5:**
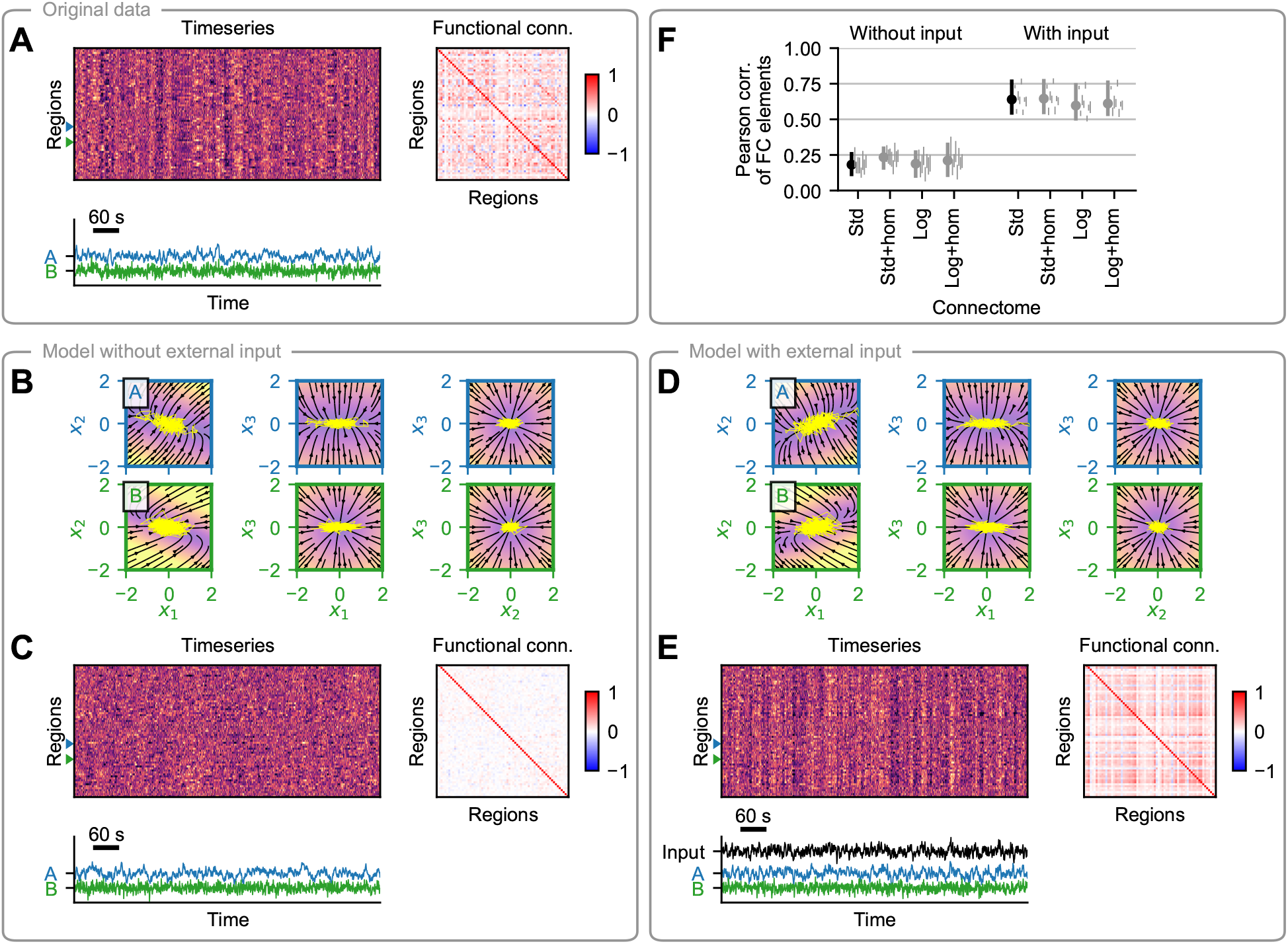
Comparison of the models for empirical HCP data. (A) Resting-state fMRI data of a single subject (top left), corresponding functional connectivity (top right), and time series of two example regions (bottom). (B) Phase plane of the inferred dynamical system without the external input. Each panel shows a two-dimensional projection of a three-dimensional system, with the third variable set to zero. The vector field is evaluated assuming zero network input and using the inferred region- and subject-specific parameters. Background color represents the velocity magnitude, yellow lines are exemplary simulated time-series of the node activity when embedded in a network. (C) Data generated by the inferred model without the external input. The regional and subject parameters are set to those inferred, while the system and observation noise is random. Layout is the same as in (A). (D-E) Same as (B-C), but for the model with the external input. The external input in (E) is generated randomly using the inferred parameters. (F) Fit between the original data and the data generated by the inferred model for all subjects. The fit is evaluated in terms of similarity of functional connectivity, measured by the Pearson correlation of the FC elements. Results are shown for the variants without and with external input, and for four variants of structural connectomes: standard (Std), standard with homotopic connections added (Std+hom), log-scaled (Log), and log-scaled with homotopic connections added (Log+hom). All lines show the [5, 95] percentile of the correlation coefficient (obtained by repeated simulations with different noise). Leftmost line in each group is the cumulative result for all subjects (with the dot showing the median value), the thin lines correspond to individual subjects. The highlighted black lines correspond to the model variants shown in (B,C) and (D,E).

As with the synthetic data, we have evaluated the goodness-of-fit by simulating new data with the trained model, and comparing their features to the original data on which the model was trained. The example of a single subject shows that while the generated time series of single regions are qualitatively similar for both the model without and with external input (Fig. 5C,E), the model with external input shows better match when considering the whole-brain features and the functional connectivity in particular. This applies for all subjects as shown by the group level quantitative analysis (Fig. 5F). The preprocessing of the structural connectome, on the other hand, has only minor effects on the goodness-of-fit (Fig. 5F). Based on these results, we focus only on the model with the external input and with standard structural connectome in the remainder of the work.

#### Two-dimensional state space sufficient for the discovered dynamics

Next, we investigate the dimensionality of the state space. An illustrative example is shown on Fig. S6, where observation of a single region is projected into the space space using models with state space dimension *n*_*s*_ from 1 to 5. No matter what the dimension of the state space of the model is, no information is encoded in the third and higher dimensions.

The same behavior is observed for all subjects and all regions in the dataset. We quantify this in Tab. 1, upper, by calculating the mean KL divergence between the instantaneous state space distribution and the cumulative state space distribution, that is of the sum of all instantaneous distributions. The results show for dimension *n*_*s*_ ≥ 2 the amount of information stored in the first and second dimension is almost constant, while virtually no information is stored in any additional dimensions.

**Table 1:** Information encoded into the dimensions of state space (top), regional parameter space (middle), and subject parameter space (bottom) when the number of dimensions is varied. In each subtable we applied the model with varying state space size *n*_*s*_, or number of regional parameter *m*_*r*_, or number of subject parameters *m*_*s*_ (columns), and then measured the information encoded in each dimension (rows). The information is quantified by the mean KL divergence between the inferred distribution (for all states at all times/all regional parameters/all subject parameters) and the cumulative distribution. By cumulative distribution we mean the sum of all state distributions for a single region (for state space), the sum of all parameter distributions for a single region (for regional parameters), or the sum of all parameter distributions (for subject parameters). The mean is calculated for every dimension of the state/parameter space separately. Since the inference process does not guarantee any specific order of state space dimensions, we order the dimensions by the amount of information stored for each model.

| State space | $n_s$ | 1 | 2 | 3 | 4 | 5 |
| --- | --- | --- | --- | --- | --- | --- |
| | $x_1$ | 2.539 | 3.252 | 3.178 | 3.195 | 3.244 |
| | $x_2$ | - | 0.383 | 0.368 | 0.378 | 0.377 |
| | $x_3$ | - | - | 0.000 | 0.000 | 0.001 |
| | $x_4$ | - | - | - | 0.000 | 0.000 |
| | $x_5$ | - | - | - | - | 0.000 |
| Regional parameters | $m_r$ | 1 | 2 | 3 | 4 | 5 |
| | $\theta_1^r$ | 2.911 | 2.767 | 2.731 | 2.731 | 2.719 |
| | $\theta_2^r$ | - | 1.958 | 1.930 | 1.949 | 1.935 |
| | $\theta_3^r$ | - | - | 0.789 | 0.836 | 0.001 |
| | $\theta_4^r$ | - | - | - | 0.000 | 0.000 |
| | $\theta_5^r$ | - | - | - | - | 0.000 |
| Subject parameters | $m_{sub}$ | 1 | 2 | 3 | 4 | 5 |
| | $\theta_1^s$ | 3.757 | 3.727 | 3.668 | 6.102 | 4.598 |
| | $\theta_2^s$ | - | 1.991 | 1.962 | 2.786 | 2.042 |
| | $\theta_3^s$ | - | - | 1.441 | 1.380 | 1.347 |
| | $\theta_4^s$ | - | - | - | 1.288 | 1.292 |
| | $\theta_5^s$ | - | - | - | - | 0.128 |

#### Three region-specific parameters identified

We can apply the same approach towards the dimensionality of region-specific parameters *m*_*r*_ as we did for the state space. Tab. 1, middle, shows the amount of information encoded in the dimensions of the parameter space as we increase the number of regional parameters. The results show that our method can identify three region-specific parameters, and adding other parameter dimensions is inutile. The amount of information encoded in the first three dimensions is stable; only when *m*_*r*_ = 5 only the first two parameters are identified, presumably due to convergence issues of the optimization problem.

What is the role of the parameters in the inferred model? We demonstrate its function by generating new data with the trained model, varying a single parameter while keeping others constant and using the same noise instantiation (Fig. 6). The first parameter 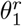 influences the presence of low-frequency (below 0.1 Hz) oscillations (Fig. 6A). The second parameter 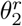 modulates the response to the external input: the simulated time series change from being correlated with the external input for negative values to anti-correlated for positive values (Fig. 6B). Finally, the third parameter 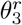 changes the response to the input from the rest of the network, from non-correlated for negative values to correlated for positive values (Fig. 6C).

**Figure 6:**
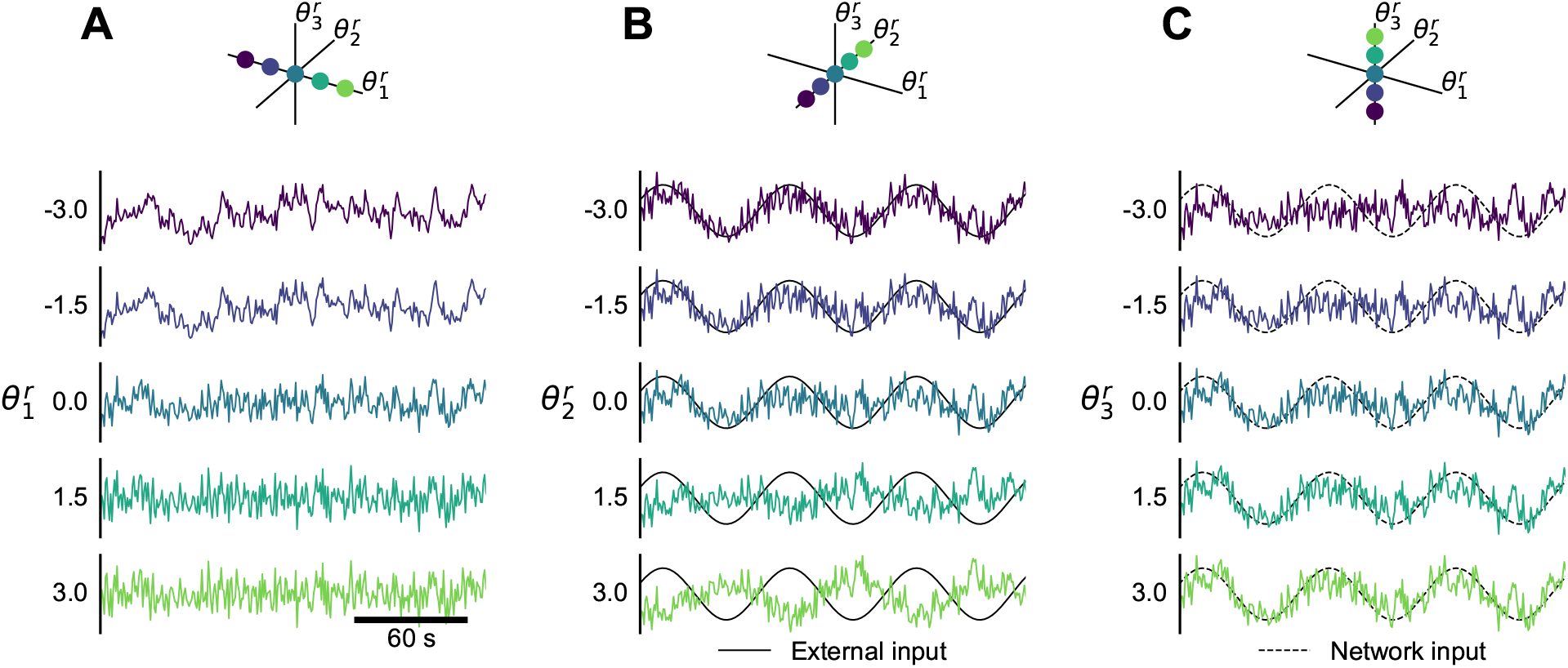
Effect of the regional parameters on the generated data. With the trained model we simulate the time series of a single region while varying one regional parameter. In each column, we systematically vary one of the region-specific parameters, while keeping others set to zero. The subject specific parameters are also set to zero. The system and observation noise are generated randomly from standard normal distributions, but kept the same for all simulations. In (A) the external input and network input are set to zero. In (B) the external input is set to a sine wave with frequency 0.014 Hz and the network input is set to zero. In (C) the external input is set to zero and the network input is set to a sine wave with frequency 0.014 Hz.

Another way of analyzing the roles of region-specific parameter is to look at their relations with various features of the structural and functional data. We divide the features in two categories: First, those derived from the individual data that were used for the model training, and second, those obtained from external sources and thus not specific to the subject in question. Taking the features and using the inferred parameters for all subjects, we performed a multivariate linear regression for the different features (Tab. 2).

**Table 2:** Results of the multivariate linear regression between the means of the inferred regional parameters ***θ***^***r***^ and regional features on the individual or population level. Shown is the coefficient of determination *R*^2^ and absolute values of weights. For visual orientation, highlighted in bold are the values where *R*^2^ *>* 0.3 and the weight *>* 0.2. In gray are weights with *p >* 0.05. The individual features are calculated from the structural connectivity (SC) or from the processed and parcellated fMRI. External data include average neuronal size and neuronal density, principal gradient of resting state functional connectivity, T1w/T2w ratio, first principal component of gene expression map, and excitation-inhibition map.

| | Feature | $R^2$ | Weights | | |
| --- | --- | --- | --- | --- | --- |
| | | | $\theta_1^r$ | $\theta_2^r$ | $\theta_3^r$ |
| Individual data | SC: Node in-strength | **0.49** | **-0.62** | -0.04 | **-0.39** |
|  | SC: Node centrality | **0.30** | **-0.46** | -0.11 | **-0.33** |
|  | fMRI: First PCA eigenvector | **0.76** | -0.20 | **-0.88** | 0.05 |
|  | fMRI: Second PCA eigenvector | 0.04 | 0.02 | 0.18 | 0.16 |
|  | fMRI: Correlation with mean signal | **0.34** | -0.14 | **-0.59** | 0.08 |
|  | fMRI: Correlation with network input | **0.41** | **-0.53** | **-0.23** | **0.36** |
|  | fMRI: Number of zero-crossings | **0.95** | **0.98** | -0.10 | -0.02 |
|  | fMRI: Power below 0.1 Hz | **0.90** | **-0.95** | 0.12 | 0.01 |
| External data | Neuronal size (Von Economo) | 0.20 | 0.40 | -0.16 | -0.19 |
|  | Neuronal density (Von Economo) | 0.27 | -0.46 | 0.21 | 0.19 |
|  | Neuronal density (BigBrain) | 0.21 | -0.41 | 0.23 | 0.04 |
|  | RSFC principal gradient | 0.10 | 0.29 | -0.16 | 0.07 |
|  | T1w/T2w ratio | 0.25 | -0.42 | 0.15 | 0.28 |
|  | Gene expression map (first PC) | **0.53** | **-0.70** | 0.19 | **0.21** |
|  | EI map | 0.11 | -0.32 | 0.10 | -0.03 |

With the individual data we evaluated the link to features of the structural connectome and of the regional fMRI time series. The results correspond well with the effects of the parameters as established above. The first parameter 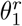 is strongly linked with the frequency features of the regional fMRI time series, specifically, the power below 0.1 Hz and number of zero-crossing. The second parameter 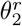 is most importantly linked with the first eigenvector obtained from principal component analysis (PCA) of the subject fMRI data. That is consistent with the interpretation that the first principal component corresponds to the external input, the response to which 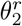 modulates. The third parameter 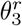 is linked to the features of the structural network, mainly node in-strength, but also to the correlation of the regional fMRI time series with the network input to the same region; this network input also depends on the structural network (Eq. 7). This is also consistent with the modulating effect of 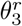 as established before. The relations between the inferred parameters and the individual data features are visualized on Fig. S7.

With the external data we compared our inferred parameters against multiple feature maps used in previous studies on regional heterogeneity in whole-brain networks (5, 6): neuronal size density and density by Von Economo and Koskinas (23), neuronal density obtained from BigBrain (24), principal gradient of resting-state functional connectivity obtained through diffusion embedding (25), T1w/T2w ratio approximating the myelin content obtained from Human Connectome Project cohort data (26), the dominant component of brain-related gene expressions (5) and excitation-inhibition (EI) map (5) obtained from Allen Human Brain Atlas (27). Only the first of our inferred parameters is strongly linked to any of them, most importantly to the gene expression map, and to lesser degree to the neuronal density from both sources, and to T1w/T2w ratio. The relations between the inferred parameters and the individual data features are visualized on Fig. S8.

#### Subject-specific parameters

Finally, we consider the role of subject-specific parameters ***θ***^*s*^. As for the regional parameters, we evaluated how much information is stored in each dimension of subject-specific parameters as we increase the number of parameters (Tab. 1, bottom). Unlike for the state space and regional parameter space, we see that the amount of information does not stabilize. Considering that even three-dimensional parameter space should be more enough to store the information about eight subjects, and also the fact that the inferred parameters do not fill the parameter space and are instead centered around the origin (Fig. 7A), we conclude that the optimization method was not able to converge properly. Therefore we need to be careful with the interpretation of the results.

**Figure 7:**
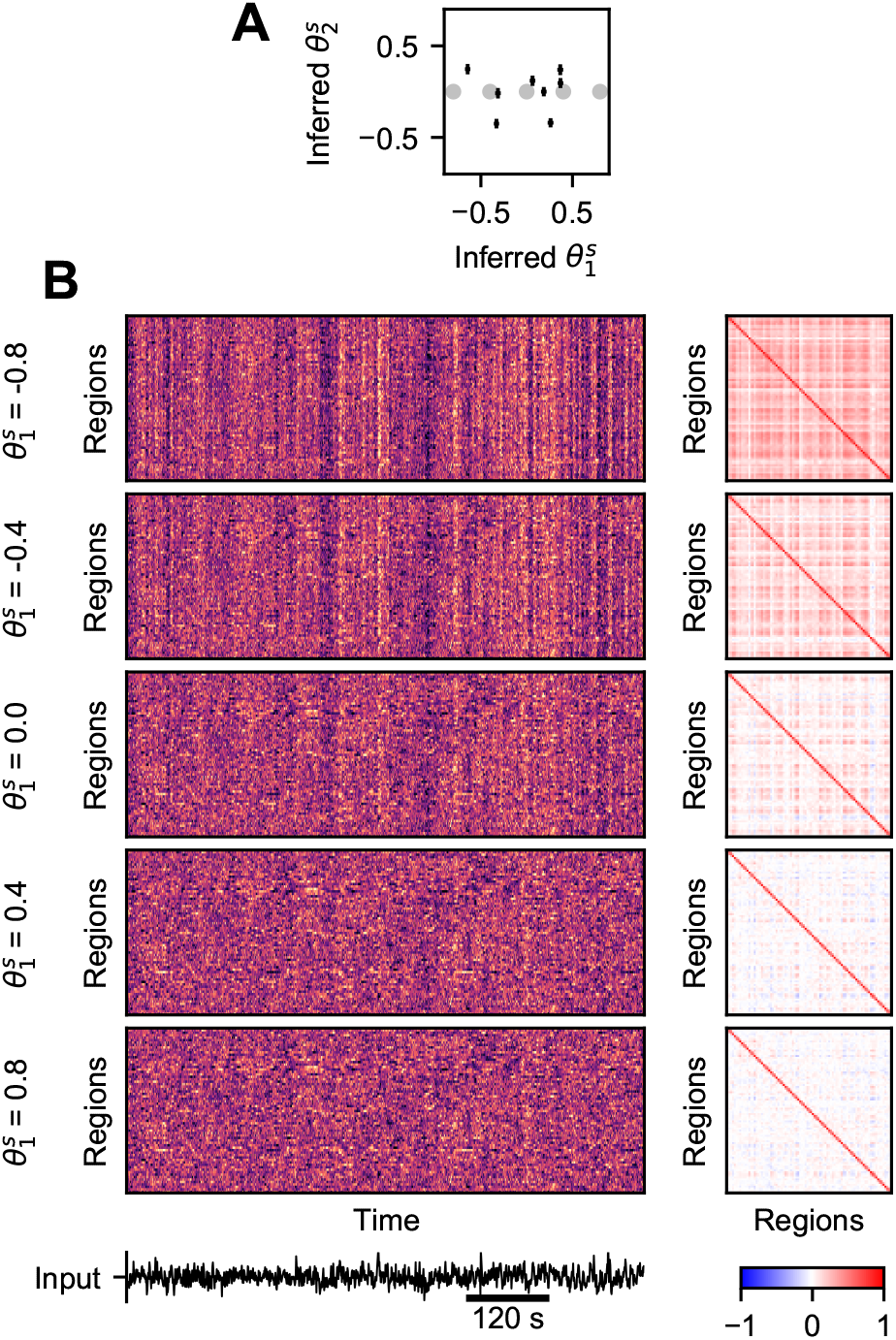
Effect of the subject-specific parameter 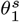 on generated whole-brain dynamics. (A) Black crosses: Inferred parameters for eight subjects in the study. Crosses indicate the inferred mean and one standard deviation. Grey dots: Parameter values for simulations in (B). (B) Data generated by the trained model, while varying the first subject-parameter. Left: Simulated time series, right: corresponding functional connectivity. Region-specific parameters are generated randomly from *N* (0, 1), but kept constant for all simulations. System and observation noise as well as the external input are also generated randomly using the inferred parameters, and kept constant between the simulations.

Nevertheless, we can still evaluate the role of the parameters by performing whole-brain simulations with the trained model while varying the parameter in question. This approach reveals that the first parameter globally modulates the effect of the external input, leading to increased (or decreased) functional connectivity (Fig. 7). It thus have a similar effect as the second regional parameter, except on whole-brain level. The second parameter (Fig. S9) however does not have a clear effect on the generated activity, apart from minor enhancement of time series maxima or minima.

Considering the low number of subjects in this study, we do try to link these subject parameter to any subject-related feature in process analogous to the one performed for regional parameters. Such approach however is possible and can reveal more about the subject-specific parameters when the cohort is sufficiently large.

## 3. Discussion

### Summary

In this work we have introduced a method for analysis of whole-brain dynamics based on a model of networked dynamical systems. Using the structure of the network and the functional data, the method allows to infer both the unknown generative dynamical system, and the parameters varying across regions and subjects.

We have tested the method on two synthetic data sets, one generated by a virtual brain model composed of nodes with a Hopf model (Fig. 2) and one generated by virtual brains with a parametric mean field model (Fig. 3). Detailed analysis of the results has shown that the proposed method can recover the original parameters as well as reproduce the important features of the original data both the single region level and on the network level (Fig. 4).

We then applied the method on human resting-state fMRI. We found that to obtain the best fit, the model needed at least two-dimensional state space and three-dimensional regional parameter space. Simulations with the inferred model revealed the distinct role of the three regional parameters: strength of low-frequency oscillations, modulation of the response to the external input, and modulation of the response to the network input (Fig. 6). The subject-specific parameter modulates the response to the external input globally (Fig. 7).

### Cortical gradients in large scale brain modeling

Gradients of structure, connectivity, gene expression, and function across the human cortex raised considerable interest in recent year (28, 29), and several modeling studies investigated their role in the large scale brain dynamics (3–6). Our approach shares with them the basic principles of large-scale brain modeling based on the structural connectome, but differs in three key points: First, we do not rely on any particular neural mass model, rather, the model is derived from data. Second, the regional parameter that we infer are not constrained by any particular parameterization. Third, via the cost function we are fitting the time series directly, and not any derived feature such as static or dynamical functional connectivity.

How do our results compare to those of previous studies? Demirtaş et al. (3) used two-population neural mass model embedded in the whole-brain network. They parameterized the neural mass parameters with the T1w/T2w ratio approximating the myelin content, and optimized the model parameter to obtain best functional connectivity fit. They found that the heterogeneity following the T1w/T2w gradient improves the fit compared to the homogeneous model or random surrogates. Wang et al. (6) used one equation neural mass model in the network, whose parameters were optimized freely in order to, again, fit the functional connectivity matrix. They found that the optimal parameters are correlated with the first principal gradient of resting-state functional connectivity (RSFC) and with the T1w/T2w map. Furthermore, the optimal strength of recurrent connection was correlated with the neuronal density map. Expanding the work, Kong et al. (4) parameterized the same model with the T1w/T2w map and RSFC gradient in order to fit not only the static, but also dynamical functional connectivity. For this goal, parameterization with both maps was necessary. These conclusions are in general agreement with our results: The first inferred parameter is correlated with T1w/T2w map, RSFC gradient, as well as neuronal density, although the relation is not as strong, particularly for the RSFC gradient (*R*^2^ = 0.10 in our case).

Kong et al. (4) however also noted that their optimal regional parameters are strongly linked with the gene expression gradients, in particular, with the first principal component of expression maps of 2413 brain-specific genes. This is a relation that our study strongly supports, as the first inferred parameter was strongly linked to the gene expression map (*R*^2^ = 0.53). Also Deco et al. (5) investigated the role of gene expression in whole-brain dynamics. They compared the goodness of fit when the neural mass model was parameterized with excitation-inhibition ratio (E:I) obtained from gene expression, as well as with first principal component of gene expression, T1w/T2w ratio, or using a homogeneous model. The fit was best in the E:I case, followed by the other heterogeneous models and then the homogeneous one. This result is not supported by this study, as the E:I map was correlated to the inferred parameters only weakly.

We have chosen to perform the analysis using the Desikan-Killiany cortical parcellation (19), mainly for consistency with related studies (4–6). It is well recognized that the choice of brain atlas can affect the results of neuroscientific studies (30). Future applications of the proposed framework might thus benefit from analyses on not only anatomy-based parcellations such as Desikan-Killiany, but also multimodal (26) or cytoarchitectonic (31) parcellations, which should better reflect the underlying structural organization.

### Learning complex dynamics

We have tested the method on synthetic data generated using the Hopf model and parametric mean field model as neural masses embedded in a whole-brain network. Admittedly, these two models, while often used in whole-brain modeling, are dynamically quite simple - after all, they are represented by one or two differential equations per node. And even for these models some shortcomings of the method are noticeable, in particular the insufficiently captured network interactions leading to weakened functional connectivity in the generated data.

One can ask whether the method would be able to handle more complex dynamics, generated by higher dimensional models, with many coexisting fixed points and limit cycles, and possibly acting on multiple time scales. In principle, the present method can be applied to more complex data, and, if the state space is set to be sufficiently large and the hidden layer in function *f* sufficiently wide, arbitrarily complex dynamics can be represented by the architecture. Whether such system can be successfully discovered through the optimization process is however a different question, one that we are not able to answer here, since designing neural network architectures for novel tasks is notoriously difficult problem without robust theoretical guidelines.

Considerable amount of other architectures were explored in related works, and although they were not applied in a networked setting, their elements could be incorporated in our framework to improve its performance for more complex dynamics. For instance, Duncker et al. (10) relied on Gaussian processes conditioned on set of fixed points to learn the system dynamics, and demonstrated its efficacy on multistable dynamical systems. Nassar et al. (12) used a tree structure to partition the state space and approximate the system in each partition with linear dynamics. Koppe et al. (15) used piecewise linear recurrent neural network to analyze fMRI data. Schmidt et al. (13) later expanded on this work introducing an approach for better approximation of systems with multiple time scales through creation of slow manifolds in the state space using a regularization scheme of the dynamical system.

### Imperfect connectome

Our method assumes that the structural connectome through which the local dynamics is coupled is known. What we can obtain, however, is only an estimate from diffusion tractography, suffering from a range of biases (32, 33). Our results indicate that while the method can handle small perturbations of the connectome, larger perturbations or different scaling can considerably degrade its performance (Fig. S5). To some extent this might be overcome by running the method with several connectomes using different scalings (linear, logarithmic) or different corrections for known biases and choosing the optimal connectome via model comparison methods.

If that would not produce results of sufficient quality, alternate approach can be pursued, one that would use the estimated structural connectome not as hard data but only as a soft prior for the effective connectivity of the model. Such approach was described for whole-brain dynamics generated by the multivariate Ornstein-Uhlenbeck process, using the thresholded structural connectivity as a topological mask for the inferred effective connectivity (34, 35). The model connectivity may be inferred even without any prior anatomical constraints, as demonstrated by the MINDy method that relies on a simple one-equation neural mass model (16).

### Importance of dynamically-relevant parameters

Large-scale brain dynamics during resting-state is altered in neurodegenerative diseases (36) and in normal aging (37). Myriads of regionally varying parameters that can plausibly influence the large-scale dynamics can be measured either in vivo or post mortem, such as cell density, cell type composition, local connectivity structure, connectivity to subcortical structures, or receptor densities, to name just a few. But which ones are in fact relevant for large-scale brain dynamics, and how do they influence it? Construction of bottom-up mechanistic models that would include all possible parameters and allow to investigate their role is unfeasible due to the complexity of human brain with its dynamics spanning multiple temporal and spatial scales, even if the parameters were in fact accurately measured (38).

Our approach instead pursues this understanding from the opposite direction. We use the amortized inference framework to learn the dynamical system driving the dynamics, and with it also the parameters varying across regions and subjects. Since these parameters are inferred from the functional data in unsupervised fashion, they are by construction the parameters relevant for the large-scale dynamics. Given the abstract nature of the inferred model, the mechanistic meaning of these dynamically-relevant parameters is not self-evident, yet they still provide a measure of similarity of brain regions and different subjects and their effect on the dynamics can be investigated through the trained model. Furthermore, given large enough data set, the dynamically-relevant parameters may be linked to the measured quantities (or their combinations). Such link may provide insights into the origin of neurodegenerative diseases if the dynamically-relevant parameters differ between the disease stages.

Importantly, the link between dynamically-relevant parameters and the measurable quantities can be estimated from a preexisting patient cohort, and then only applied to single subject. That is advantageous if the measurement is difficult, costly, or impossible to perform in clinical setting (such as for cell type composition estimated from post mortem studies); in such cases, the dynamically-relevant parameters may instead be estimated from easy-to-obtain resting-state fMRI and then mapped using the known link. This approach thus opens new possibilities for exploitation of large scale neuroimaging databases such as Human Connectome Project (18) or UK Biobank (39) on one hand and detailed cytoarchitectonic (24, 31) or genetic (27) brain atlases on the other.

## 4. Methods

### 4.1. Structural connectomes

The structural connectomes used for the application on empirical data, as well as for the generation of synthetic data sets, were derived from the neuroimaging data from Human Connectome Project (18). Specifically, eight random subjects from HCP 1200 Subjects cohort were used (ID numbers 100307, 100408, 101107, 101309, 101915, 103111, 103414, and 103818). For those, *Structural Preprocessed* and *Diffusion Preprocessed* packages were downloaded (40). Next, the structural connectomes were built for the cortical regions of Desikan-Killiany parcellation (19) using MRtrix 3.0 (41). To do so, first the response function for spherical deconvolution was estimated using the *dhollander* algorithm (42). Next, fibre orientation distribution was estimated using multi-shell multi-tissue constrained spherical deconvolution (43). Then 10 million tracks were generated using the probabilistic iFOD2 (second-order integration over fiber orientation distributions) algorithm (44). These were then filtered using the SIFT algorithm (45). Finally, the connectome were built by counting the tracks connecting all pairs of brain regions in the parcellation.

Four variants of the structural connectomes were used for the application on empirical data: The standard variant corresponds to the description above. The log-scaled connectome was calculated as *W*_log_ = log_10_(*W* + 10^*q*^) with *q* = −3, *W* being the original connectome. Both the standard and log-scaled connectome were also modified by strengthening the homotopic connections, that is those connecting corresponding regions in the opposite hemispheres. For this, the strength of all homotopic connections were set to 97-percentile of all values in the structural connectome. The perturbed connectomes were constructed by taking the original connectome *W* and adding a matrix with elements from random normal distribution, scaled by the perturbation magnitude *ϵ*, i.e. *W*_*ϵ*_ = *W* + *ϵA*. For each value of perturbation magnitude, four different perturbed connectomes were built. In all cases, the connectome matrices were normalized so that the largest element in each was equal to one.

### 4.2. Resting-state fMRI data

The resting-state fMRI data were obtained from the Human Connectome Project for the same eight subjects as the structural data. We have used the resting-state data preprocessed by the HCP functional pipeline and ICA-FIX pipeline (*Resting State fMRI FIX-Denoised* package). These were further processed by the DiCER method (22) with default parameters. The DiCER method was designed to remove widespread deflection from the fMRI data, and provide better alternative to global signal regression. Afterwards, the processed data were parcellated into 68 regions of Desikan-Killiany parcellation (19), and each time series was normalized to zero mean and unit variance. The DiCER preprocessing was performed for all four sessions (REST1_LR, REST1_RL, REST2_LR, REST2_RL) concatenated. However only the results from the first session (REST1_LR) were utilized in the study. The time series were thus 14.4 minutes long (1200 samples with 0.72 Hz sampling frequency).

### 4.3. Amortized variational inference for networks of nonlinear dynamical systems

#### Generative dynamical system

As outlined above, we assume that the observed activity *y*_*j*_(*t*) of a brain region *j* is generated by a dynamical system,

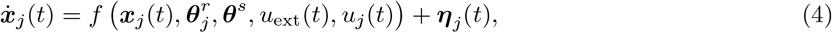

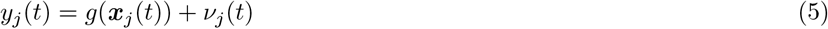

where 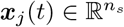 is the state at time 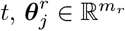 and 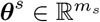 are the region-specific and subject-specific parameters, *u*_ext_(*t*) is the external input, shared by all regions of a single subject,

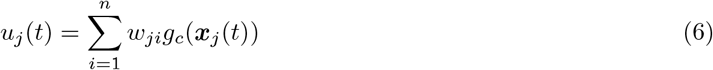

is the network input to region *j* with 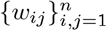 being the structural connectome matrix of the network with *n* nodes.

To make the inference problem more tractable, we simplify the problem and assume that the nodes are coupled through the observed variable *y*_*j*_. More precisely, we assume that in Eqs. (4-6) *g* ≡ *g*_*c*_, and that the observation noise term *ν*_*j*_ is small enough that it can be included in the coupling. Then the network input has the form

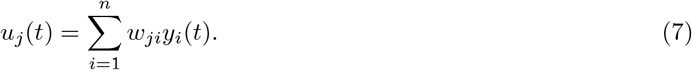

This form has the advantage that the network input is independent of any hidden variables and can be computed directly from the known observations *y*_*j*_. This effectively decouples the time series in different nodes so that they can be processed separately, as described below.

For the purpose of the inference, we use the time-discretized form of Eqs. (4-5) utilizing the Euler method,

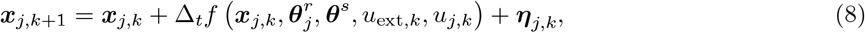

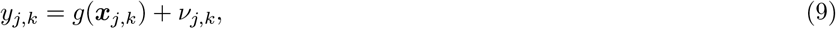

where we denote the time step with the index *k*.

*Evidence lower bound*. As usual in variational inference, we aim to maximize the evidence lower bound (ELBO), and by doing so at the same time minimize the Kullback-Leibler divergence between the true posterior and the approximate posterior *q*. In the following text, we consider only a single data point from one subject and one region, and we omit the region indexing for brevity.

A single data point {***y, u, c***} representing the data from a one region is composed of the observed time series 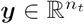, network input time series 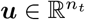, and one-hot vector 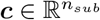, that is, a vector with zeros everywhere except *i*-th position with value one, encoding the identity of subject *i*. For this data point the ELBO can be expressed as follows. (For details see Supplementary Information.)

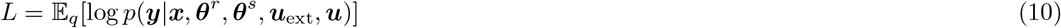

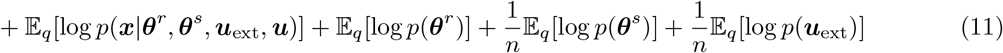

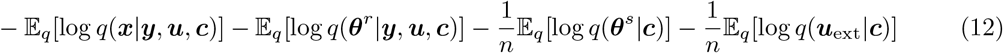

Here the first line represents the decoder loss, second line the priors for states ***x***, region- and subject-specific parameters ***θ***^*r*^ and ***θ***^*s*^, and the external input ***u***_ext_, and the third line the approximate posteriors again for states, region-, subject-specific parameters, and the external input.

#### Decoder, or the observation model

We assume that the observation model can be modeled as a linear transformation of the system state with Gaussian noise, *y* = *g*(***x***) + *ν* = ***a***·***x*** + *b* + *ν*. This forward projection essentially represents the decoder part of the encoder-decoder system, and so the likelihood in Eq. (10) can be expanded over time as

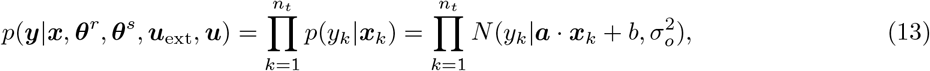

where *N* (*y*|*μ, σ*^2^) stands for normal distribution with mean *μ* and variance *σ*^2^. The parameters of the observation model, which are to be optimized, are the coefficients of the linear projection ***a*** and *b*, together with the observation noise variance 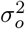.

#### Prior on the system states

The first term in Eq. (11) represents the prior function on the system states ***x*** given the network input ***u***, external input ***u***_ext_, and the parameters ***θ***^*r*^ and ***θ***^*s*^. It is here where the dynamical system *f* appears in the ELBO. This term can be expanded over time as

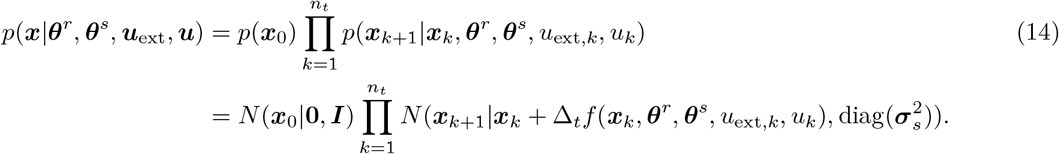

Here we use the standard normal distribution as a prior for the initial state ***x***_0_, and then evolve the system over time according to the function *f*. We represent the function *f* as a two-layer neural network, with a Rectified Linear Unit (ReLU) activation function in the hidden layer. The weights of the network are to be optimized, together with the system noise standard deviation ***σ***_*s*_. The number of hidden units is given in the Tab. 3.

**Table 3:**
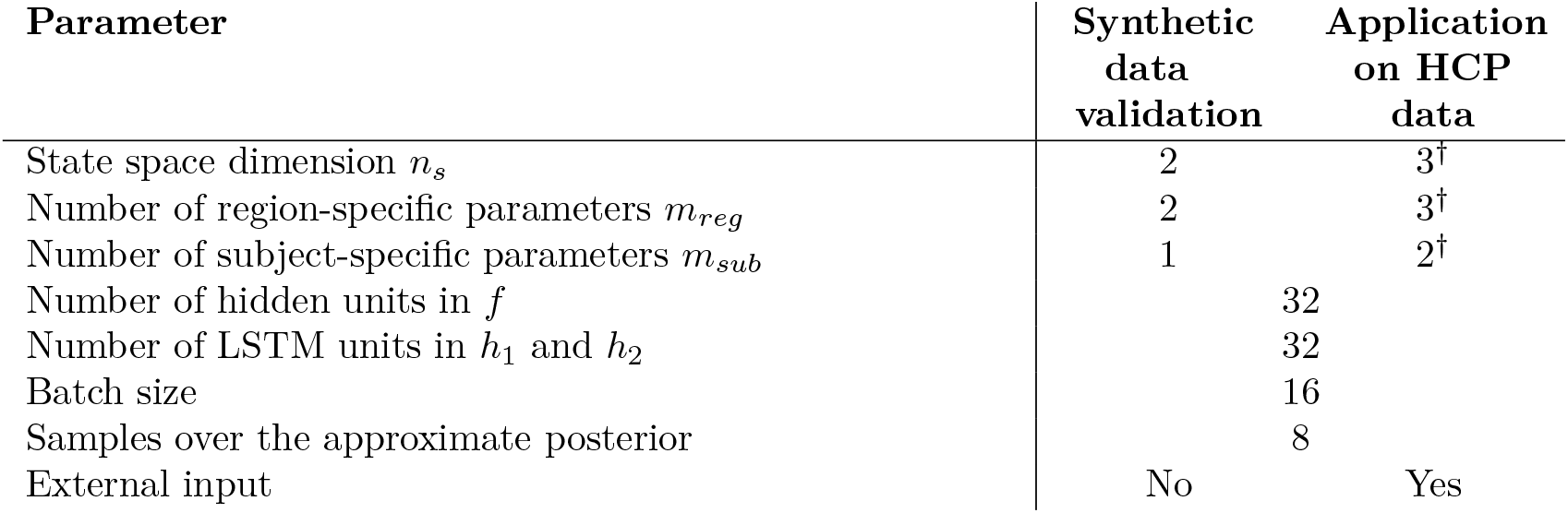
Method parameters used in the test cases on synthetic data and for the application on HCP resting-state fMRI data. ^†^Or varying.

#### Prior on the parameters

For the region- and subject-specific parameters we utilize the standard normal distribution as a prior, as is often used in variational autoencoders. The priors in the second and the third term in Eq. (11) can thus be written as *p*(***θ***^*r*^) = *N* (***θ***^*r*^|**0, *I***) and *p*(***θ***^*s*^) = *N* (***θ***^*s*^|**0, *I***).

#### Prior on the external input

We set the prior for the external input to an autoregressive process with time scale *τ* and variance *σ*^2^. Then the prior reads

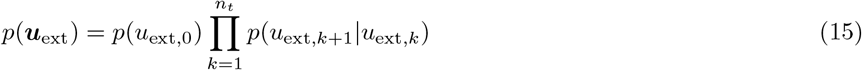

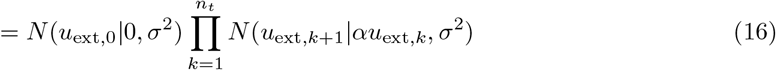

with *α* = *e*^−1*/τ*^. The variance is fixed to *σ* = 1, since any scaling can be done inside the function *f*, and the time scale is optimized together with the other parameters and neural network weights in the optimization process.

#### Approximate posteriors

We follow the standard approach and utilize multivariate normal distributions for the approximate posteriors in Eq. (12). For the states ***x*** and region-specific parameters ***θ***^*r*^ we use the idea of amortized variational inference and instead of representing the parameters directly, we train a recurrent neural network to extract the means and the variances from the time series of the observations ***y***, time series of the network input ***u***, and the one-hot vector ***c*** encoding the subject identity:

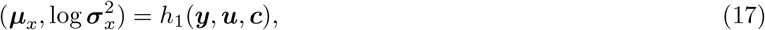

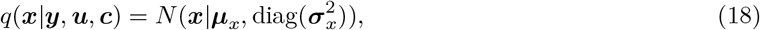

and

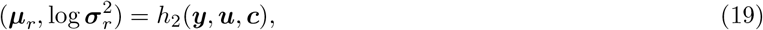

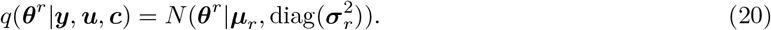

Specifically, we use Long Short-Term Memory (LSTM) networks for both functions *h*_1_ and *h*_2_. The input to the networks at step *k* is the concatenated observation *y*_*k*_ and the network input *u*_*k*_, to which is also appended the time-independent one-hot vector ***c***.

The subject-specific parameters ***θ***^*s*^ and the external input ***u***_ext_ depend only on the subject identity encoded in the one-hot vector ***c***. Their means and variances are stored directly in the matrices of means (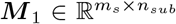for ***θ***^*s*^ and 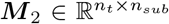 for ***u***_ext_) and matrices of log-variances (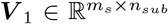 and 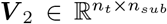). For a specific subject the relevant values are extracted through the product with the one-hot vector ***c***,

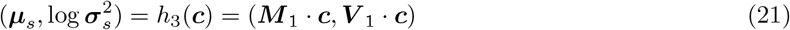

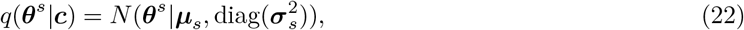

and

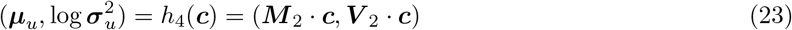

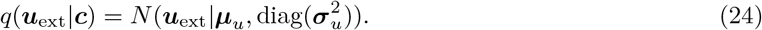

#### Optimization

The optimization target is the negative dataset ELBO,

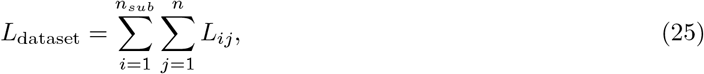

where *L*_*ij*_ is the ELBO associated with a subject *i* and region *j*, defined by Eqs. (10-12). We minimize the cost function over the weights of the LSTM networks *h*_1_, *h*_2_, weights of the neural network *f*, means and variances of the subject-specific parameters and of the external input time series ***M***_1_, ***M***_2_, ***V***_1_, ***V***_2_, external input time scale *τ*, system and observation noise variances 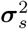 and 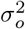 (in log-scale), and forward projection parameters ***A*** and ***b***.

The method is implemented in Keras 2.4 (46). The parameters of the method and of optimization procedure are given in Tab. 3. For optimization we use the Adam algorithm (47). The expectations in Eqs. (10-12) are approximated using samples drawn from the approximate posterior distribution. The optimization is run for 2000 epochs with learning rate 0.003 and then for additional 1000 epochs with learning rate 0.001. To make the optimization more stable we use gradient clipping with limits (−1000, 1000). To better guide the optimization procedure, we follow the previous works (14) with initial ELBO relaxation: The terms corresponding to the priors and approximate posteriors for states ***x*** and parameters ***θ***^*r*^ and ***θ***^*s*^ (Eqs. (11-12)) are scaled by a coefficient *β*, which linearly increases from 0 to 1 between the first and 500th epoch.

Two regularization terms are added to the cost function. First is a L2 regularization on the kernel weights biases and of the neural network representing function 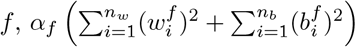, where *n*_*w*_ and *n*_*b*_ is the number of kernel weights 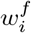 and bias coefficients 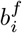, respectively. Second is on the states ***x***, 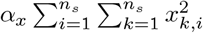. We set *α*_*f*_ = 0.01 and *α*_*x*_ = 0.01.

The initial conditions for the optimization are set as follows. The log variances of the system noise are set to −2, and the log variances of the observation noise to 0. The projection vector ***a*** is initialized randomly drawing from normal distribution (mean 0, std 0.3), and the projection bias *b* is set to zero. Matrices for subject-specific parameters and for external input ***M***_1_, ***M***_2_, ***V***_1_, ***V***_2_ are initialized randomly drawing from a normal distribution (mean 0, std 0.01). All layers of employed neural networks use the default initialization provided by Keras.

### 4.4. Whole-brain network models for simulated data sets

#### Hopf bifurcation model

The Hopf model of large-scale brain dynamics (8) is built by placing a neural mass near supercritical Hopf bifurcation at each node of a brain network. Each neural mass *i* is described by two parameters: bifurcation parameter *a*_*i*_ and intrinsic frequency *f*_*i*_. For *a*_*i*_ *<* 0 the uncoupled neural mass has one stable fixed point, and for *a*_*i*_ *>* 0 the neural mass has a stable limit cycle indicating sustained oscillations with frequency *f*_*i*_. The bifurcation exists at the critical value *a*_*i*_ = 0. The dynamics of each node in the network are given by a set of two coupled nonlinear stochastic differential equations,

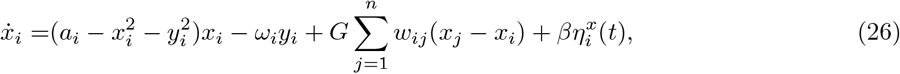

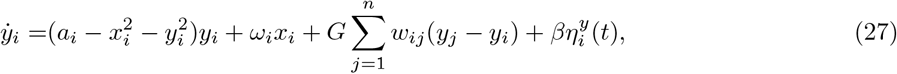

where *ω*_*i*_ = 2*πf*_*i*_, *G >* 0 is the scaling of the coupling, *w*_*ij*_ is the weight of connection from node *j* to node *i*. Additive Gaussian noise *η* is included in the equations, with standard deviation *β*.

To generate the synthetic dataset, we use the structural connectome matrices of human subjects as described above. We simulate eight subjects, with increasing coupling coefficient *G* spaced linearly between 0 and 0.7. The intrinsic frequency *f*_*i*_ of all nodes is sampled randomly from uniform distribution on [0.03, 0.07] Hz. The bifurcation parameter *a* is sampled randomly from uniform distribution [−1, 1]. The initial conditions of the system for all subjects and both variables are chosen randomly from normal distribution *N* (0, 0.3), the system is then simulated for 205 s. First 25 s are then discarded to avoid the influence of the initial conditions, leaving 180 s of data. The system is simulated with Euler method with time step Δ_*t*_ = 0.02 s. As the observed variable we take the first of the two variables in each node (i.e., *x*_*i*_), downsampled to 1 Hz, therefore every timeseries contain 180 time points. The data are normalized to zero mean and variance equal to one (calculated across the whole data set).

#### Parametric mean field model

The parametric mean field model (pMFM) was derived as a reduction from a spiking neural model (7). The resulting model is described by one nonlinear stochastic differential equation in each node of the brain network,

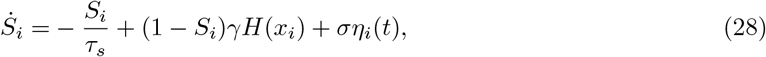

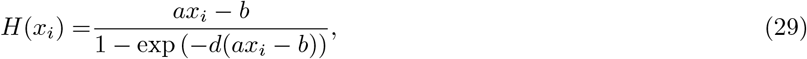

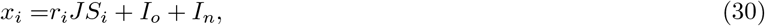

where *x*_*i*_ is the total input current, *H*(*x*_*i*_) is the population firing rate, and *S*_*i*_ the average synaptic gating variable. The total input current depends on the recurrent connection strength *r*_*i*_, synaptic coupling strength *J* = 0.2609 nA, excitatory subcortical input *I*_*o*_ = 0.295 nA, and the regional coupling 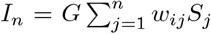, scaled by the global scaling coefficient *G*. The strength of the coupling between region *j* and *i* is proportional to the structural connection strength *w*_*ij*_. The kinetic parameters of the models are the decay time constant *τ*_*s*_ = 100 ms and *γ* = 0.641*/*1000. Values for the input-output function *H*(*x*_*i*_) are *a* = 270 nC^−1^, *b* = 108 Hz, *d* = 0.154 s. Depending on the parameter values and the strength of the network coupling, the system can be either in monostable downstate regime at low firing-rate values, bistable regime with two stable fixed points, or monostable upstate regime at high firing-rate values. The stochastic transitions between states are driven by the additive Gaussian noise *η*_*i*_ with standard deviation *σ*.

The initial conditions for *S*_*i*_ were chosen randomly from uniform distribution on [0.2, 0.8]. The system was simulated for eight subjects with connectome matrices described above. For each subject, a specific value of coupling coefficient *G* producing the strongest functional connectivity was used. This was determined by performing 4 minute long simulations with subject-specific connectome and fixed regionally heterogeneous parameters, repeated for 31 values of *G* between 0.17 and 0.22 (where optimal value was expected to lie), and picking the value where the mean of functional connectivity from the last 2 minutes was the highest. With this value of *G*, the activity of each subject was simulated for 16.4 minutes, first two of which were discarded to avoid the influence of the initial conditions. The Euler method with time step Δ_*t*_ = 10 ms was used for the simulation. The resulting time series of *S*_*i*_ were temporally averaged over windows of size 0.72 seconds, leaving 1200 time points in every time series. The data are normalized to zero mean and variance equal to one (calculated across the whole data set).

### 4.5. Relation of regional parameters and regional features from individual and external data

To analyze the role of the regional parameters inferred from human resting-state fMRI data we compare the inferred parameters to several regional features obtained on the individual level or on a population level from previous literature. All features are represented by a vector of 68 elements, corresponding to the cortical regions of Desikan-Killiany parcellation.

#### Features from individual data

The individual-level features are derived from the data used in the model fitting: structural connectivity and parcellated resting-state fMRI. For structural connectivity it is node in-strength and eigenvector centrality. For fMRI data it is the first and second spatial eigenvector obtained from principal component analysis, vector of correlation coefficients of regional time series with the mean signal of resting-state fMRI, vector of correlation coefficients of regional time series with the network input time series (Eq. 7), number of zero-crossings of the regional signals, and the power below 0.1 Hz. We note that the signals are all normalized, so the total power is constant.

#### Features from external data

We further consider several regional features derived from other sources unrelated to the modeling data. First, it is the neuronal density and neuronal size derived from the pioneering work of Von Economo and Koskinas (23). These data were mapped to Desikan-Killiany atlas (48) and used previously in related large-scale brain modeling study (6). Our inspection of these remapped data however indicated possible error in the mapping, as it contained density values that would be obtained by summing across cortical sublayers instead of averaging. Therefore we opted to redo the mapping, using the tabular data given in the recent translation of the original work (49). We followed the procedure as described in (48), only taking an average of neuronal densities and sizes where multiple sublayers were given.

In addition, we have used neuronal density estimated from the BigBrain model (24). The BigBrain model is a 3D-reconstructed data set with a spatial resolution of 20 micrometre isotropic, i.e. nearly cellular. The regional cell densities were estimated by randomly sampling at least 15 3D chunks per region from the cortex of the BigBrain model. For each region in the parcellation map, between 15 and 140 random coordinates were drawn in the ICBM 2009c nonlinear asymmetric space depending on the size of each region (50). Each coordinate was transformed to the BigBrain histological space using a nonlinear transformation guided by sulcal constraints (51), and then shifted to the closest position around the cortical mid-surface, as defined by the BigBrain layer segmentation (52). A square cube extending across the full cortical depth was sampled at each mid-surface location, and layer-specific histograms of BigBrain grayvalues were extracted from each cube. The grayvalues are 8 bit integers in the range 0… 255, where dark values indicate the presence of cell bodies due to the silver staining that was applied to the BigBrain tissue sections. To convert gray values into actual density estimates, a collection of 120 BigBrain cortical image patches scanned at 1 micron resolution (e.g. (53); full list of data references in Tab. S1) with explicit layer-specific density estimates in numbers of segmented cell bodies (54) per 0.1 cube millimeter was used to compute an individual calibration function per layer. These calibration functions were applied to the mean grayvalue per layer in each sampled cube. From the resulting layerwise densities in each sampled cube, average cortical cell densities were determined by weighting each layer density with its relative volume. The final cell density estimate for each parcellation region was determined as the average cortical density across all cubes sampled from it.

Next, the resting-state functional connectivity (RSFC) principal gradient (25) was obtained from resting-state fMRI of large cohort of healthy adults by means of diffusion embedding, a non-linear projection technique. It was proposed as a proxy measure for processing hierarchy. We use the numerical values provided by Kong et al. (4). The T1w/T2w ratio is used as a measure of myelin content. We use the map in Desikan-Killiany parcellation (4) obtained from Human Connectome Project dataset of 1200 healthy adults (26). Finally, we consider two features derived from gene expression profiles (5) from Allen Human Brain Atlas (27): First is the first principal component of 1926 brain-relevant genes. Second is the excitation-inhibition (EI) map, estimated from the expressions of genes coding for the excitatory AMPA and NMDA receptors and inhibitory GABA_A_ receptors.

#### Multivariate linear regression

All features were normalized before performing the linear regression. As the regional parameters ***θ***^*r*^ were estimated probabilistically, that is, we inferred the mean and variance of a normal distribution, we performed the linear regression on hundred samples from these inferred distributions.

## Acknowledgements

This work was supported by a public grant overseen by the French National Research Agency (ANR) as part of the second “Investissements d’Avenir” program (reference: ANR-17-RHUS-0004) and by the European Union’s Horizon 2020 research and innovation programme under grant agreement No. 945539 (SGA3) Human Brain Project. Data were provided by the Human Connectome Project, WU-Minn Consortium (Principal Investigators: David Van Essen and Kamil Ugurbil; 1U54MH091657) funded by the 16 NIH Institutes and Centers that support the NIH Blueprint for Neuroscience Research; and by the McDonnell Center for Systems Neuroscience at Washington University. The authors wish to thank Jayant Jha and Anirudh N. Vattikonda for helpful discussions.

## Data and code availability

The code associated with this study is available at https://github.com/ins-amu/SipEtAl22_ParamInferenceWithUnknownDynamics. The data are available from the portal of Human Connectome Project https://db.humanconnectome.org/.

