## Supplementary information for "Parameter inference on brain network models with unknown node dynamics and spatial heterogeneity"

### Evidence lower bound (ELBO)

For a single subject, the observations contain the time series from all $n$ regions, $\boldsymbol{Y} = (\boldsymbol{y}_1, \ldots, \boldsymbol{y}_n) \in \mathbb{R}^{n\, n_t}$, where $n_t$ is the number of time points. They are complemented by the region time series for the network input, $\boldsymbol{U} = (\boldsymbol{u}_1, \ldots, \boldsymbol{u}_n) \in \mathbb{R}^{n\, n_t}$, and the one-hot vector $\boldsymbol{c} \in \mathbb{R}^{n_{sub}}$ encoding the subject identity. The latent variables $\boldsymbol{Z}$ contain the state time series $\boldsymbol{x}_j \in \mathbb{R}^{n_s n_t}$ for all regions $j$ (where $n_s$ is the dimension of the state space), region-specific parameters $\boldsymbol{\theta}^r_j \in \mathbb{R}^{m_r}$, subject specific parameters $\boldsymbol{\theta}^s \in \mathbb{R}^{m_s}$, and time series of the external input $\boldsymbol{u}_{\text{ext}} \in \mathbb{R}^{n_t}$. The latent variables can be thus written as $\boldsymbol{Z} = (\boldsymbol{x}_1, \ldots, \boldsymbol{x}_n, \boldsymbol{\theta}^r_1, \ldots, \boldsymbol{\theta}^r_n, \boldsymbol{\theta}^s, \boldsymbol{u}_{\text{ext}})$. Our goal is to minimize the Kullback-Leibler divergence between the approximate and true posterior, which can be rewritten as a sum of subject ELBO and evidence itself,

$$\begin{aligned}
&\mathrm{KL}(q(\boldsymbol{Z}|\boldsymbol{Y},\boldsymbol{U},\boldsymbol{c}) \;||\; p(\boldsymbol{Z}|\boldsymbol{Y},\boldsymbol{U},\boldsymbol{c})) = \mathbb{E}_q[\log q(\boldsymbol{Z}|\boldsymbol{Y},\boldsymbol{U},\boldsymbol{c})] - \mathbb{E}_q[\log p(\boldsymbol{Z}|\boldsymbol{Y},\boldsymbol{U},\boldsymbol{c})] \\
&= \underbrace{\mathbb{E}_q[\log q(\boldsymbol{Z}|\boldsymbol{Y},\boldsymbol{U},\boldsymbol{c})] - \mathbb{E}_q[\log p(\boldsymbol{Y}|\boldsymbol{Z},\boldsymbol{U},\boldsymbol{c})] - \mathbb{E}_q[\log p(\boldsymbol{Z}|\boldsymbol{U},\boldsymbol{c})]}_{-L_{\text{subject}}} + \mathbb{E}_q[\log p(\boldsymbol{Y}|\boldsymbol{U},\boldsymbol{c})].
\end{aligned}$$

Maximizing the ELBO then minimizes the KL divergence. We can factorize all terms of the ELBO across $n$ brain regions: the approximate posterior,

$$q(\boldsymbol{Z}|\boldsymbol{Y},\boldsymbol{U},\boldsymbol{c}) = \prod_{j=1}^{n} q(\boldsymbol{x}_j|\boldsymbol{y}_j,\boldsymbol{u}_j,\boldsymbol{c}) \prod_{j=1}^{n} q(\boldsymbol{\theta}^r_j|\boldsymbol{y}_j,\boldsymbol{u}_j,\boldsymbol{c})\, q(\boldsymbol{\theta}^s|\boldsymbol{c})\, q(\boldsymbol{u}_{\text{ext}}|\boldsymbol{c}),$$

the data likelihood,

$$p(\boldsymbol{Y}|\boldsymbol{Z},\boldsymbol{U},\boldsymbol{c}) = \prod_{j=1}^{n} p(\boldsymbol{y}_j|\boldsymbol{x}_j,\boldsymbol{\theta}^r_j,\boldsymbol{\theta}^s,\boldsymbol{u}_{\text{ext}},\boldsymbol{u}_j,\boldsymbol{c}),$$

and the prior,

$$\begin{aligned}
p(\boldsymbol{Z}|\boldsymbol{U},\boldsymbol{c}) &= p(\boldsymbol{x}_1,\ldots,\boldsymbol{x}_n|\boldsymbol{\theta}^r_1,\ldots,\boldsymbol{\theta}^r_n,\boldsymbol{\theta}^s,\boldsymbol{u}_{\text{ext}},\boldsymbol{U},\boldsymbol{c})\; p(\boldsymbol{\theta}^r_1,\ldots,\boldsymbol{\theta}^r_n,\boldsymbol{\theta}^s,\boldsymbol{u}_{\text{ext}}|\boldsymbol{U},\boldsymbol{c}) \\
&= \prod_{j=1}^{n} p(\boldsymbol{x}_j|\boldsymbol{\theta}^r_j,\boldsymbol{\theta}^s,\boldsymbol{u}_{\text{ext}},\boldsymbol{u}_j,\boldsymbol{c}) \prod_{j=1}^{n} p(\boldsymbol{\theta}^r_j|\boldsymbol{u}_j,\boldsymbol{c})\, p(\boldsymbol{\theta}^s|\boldsymbol{c})\, p(\boldsymbol{u}_{\text{ext}}|\boldsymbol{c}).
\end{aligned}$$

We require that the data likelihood and priors depend on the subject identity only through the latent variables, so we remove the dependence on $\boldsymbol{c}$. We also require that the priors of $\boldsymbol{\theta}^r_j$ do not depend on the external input $\boldsymbol{u}_j$. Then we define the region ELBO as

$$\begin{aligned}
L_j = &\;\mathbb{E}_q[\log p(\boldsymbol{y}_j|\boldsymbol{x}_j,\boldsymbol{\theta}^r_j,\boldsymbol{\theta}^s,\boldsymbol{u}_{\text{ext}},\boldsymbol{u}_j)] \\
&+ \mathbb{E}_q[\log p(\boldsymbol{x}_j|\boldsymbol{\theta}^r_j,\boldsymbol{\theta}^s,\boldsymbol{u}_{\text{ext}},\boldsymbol{u}_j)] + \mathbb{E}_q[\log p(\boldsymbol{\theta}^r_j)] + \frac{1}{n}\mathbb{E}_q[\log p(\boldsymbol{\theta}^s)] + \frac{1}{n}\mathbb{E}_q[\log p(\boldsymbol{u}_{\text{ext}})] \\
&- \mathbb{E}_q[\log q(\boldsymbol{x}_j|\boldsymbol{y}_j,\boldsymbol{u}_j,\boldsymbol{c})] - \mathbb{E}_q[\log q(\boldsymbol{\theta}^r_j|\boldsymbol{y}_j,\boldsymbol{u}_j,\boldsymbol{c})] - \frac{1}{n}\mathbb{E}_q[\log q(\boldsymbol{\theta}^s|\boldsymbol{c})] - \frac{1}{n}\mathbb{E}_q[\log q(\boldsymbol{u}_{\text{ext}}|\boldsymbol{c})]
\end{aligned}$$

so that the subject ELBO is the sum of region ELBOs, $L_{\text{subject}} = \sum_{j=1}^{n} L_j$.

| Area | Reference |
|---|---|
| Area FG4 (FusG) | 1 |
| Area 44 (IFG) | 2 |
| Area 4a (PreCG) | 3 |
| Area 7P (SPL) | 4 |
| Area FG1 (FusG) | 5 |
| Area Fp1 (FPole) | 6 |
| Area hOc1 (V1, 17, CalcS) | 7 |
| Area hOc2 (V2, 18) | 8 |
| Area hOc3d (Cuneus) | 9 |
| Area hOc4d (Cuneus) | 10 |
| Area hOc5 (LOC) | 11 |

Table S1: Data references on EBRAINS portal for layer-specific distributions of segmented cells BigBrain model.

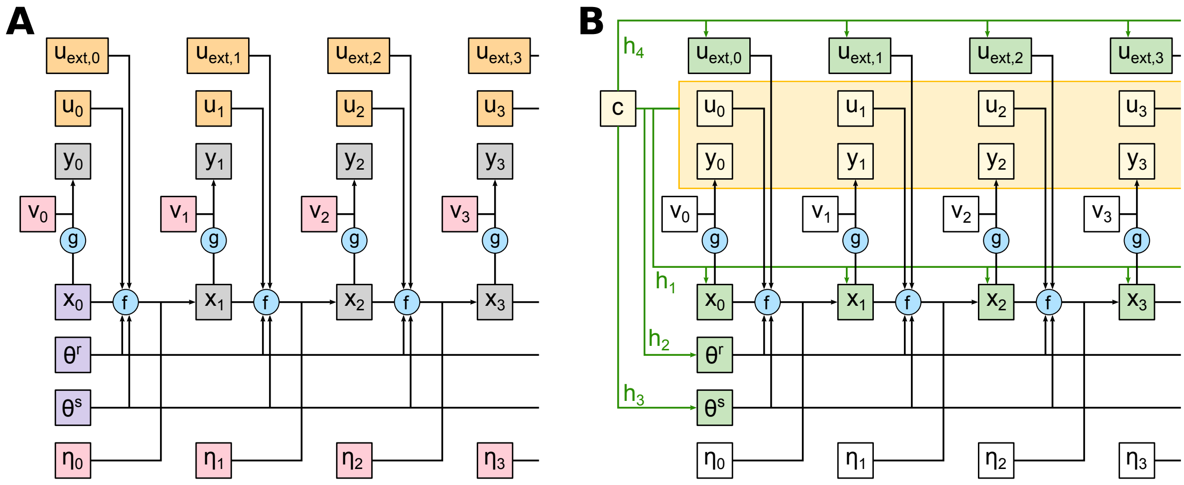


Figure S1: Overview of the method architecture visualized for one brain region. In the sketches we drop the region indices for simplicity, and keep only the time indices. (A) Generative model. With known functions $f$ and $g$, and given initial conditions $\boldsymbol{x}_0$, parameters $\boldsymbol{\theta}^r$ and $\boldsymbol{\theta}^s$ and time-varying external input $u_{\text{ext}}$, the model can be simulated in forward fashion, influenced by the system noise $\boldsymbol{\eta}$ and observation noise $\nu$. The network input for region $j$ at time $k$ is calculated on the fly from the current states of other regions, $u_{j,k} = \sum_{i=1}^{n} w_{ji} y_{i,k}$. (B) Inference model. The data (observation time series $\boldsymbol{y}$, precomputed network input time-series $\boldsymbol{u}$ and one-hot vector $\boldsymbol{c}$ identifying the subject) are mapped through the encoder functions $h_1$, $h_2$, $h_3$, and $h_4$ onto the system states $\boldsymbol{x}$, region-specific parameters $\boldsymbol{\theta}^r$, subject-specific parameters $\boldsymbol{\theta}^s$, and external input $u_{\text{ext}}$ respectively. The observation function $g$ appears in the likelihood function, while the system evolution function $f$ enters the prior on the states. The noise $\boldsymbol{\eta}$ and $\nu$ is present only implicitly via the likelihood and the prior functions. The inference problem amounts to the maximization of the resulting ELBO over the parameters of the generative model $f$, $g$, encoder functions $h_1$, $h_2$, $h_3$, $h_4$, and variance of the system and observation noise.

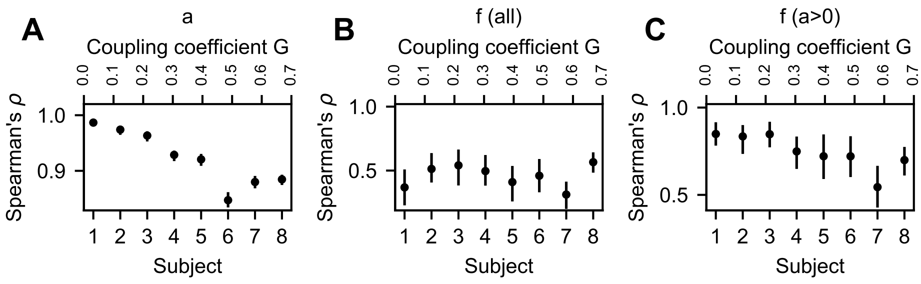


Figure S2: Recovery of the region-specific parameters in the Hopf model for different subjects with different coupling coefficient $G$. The figure contains the data from Fig. 2A in the main text, separated for the individual subjects.

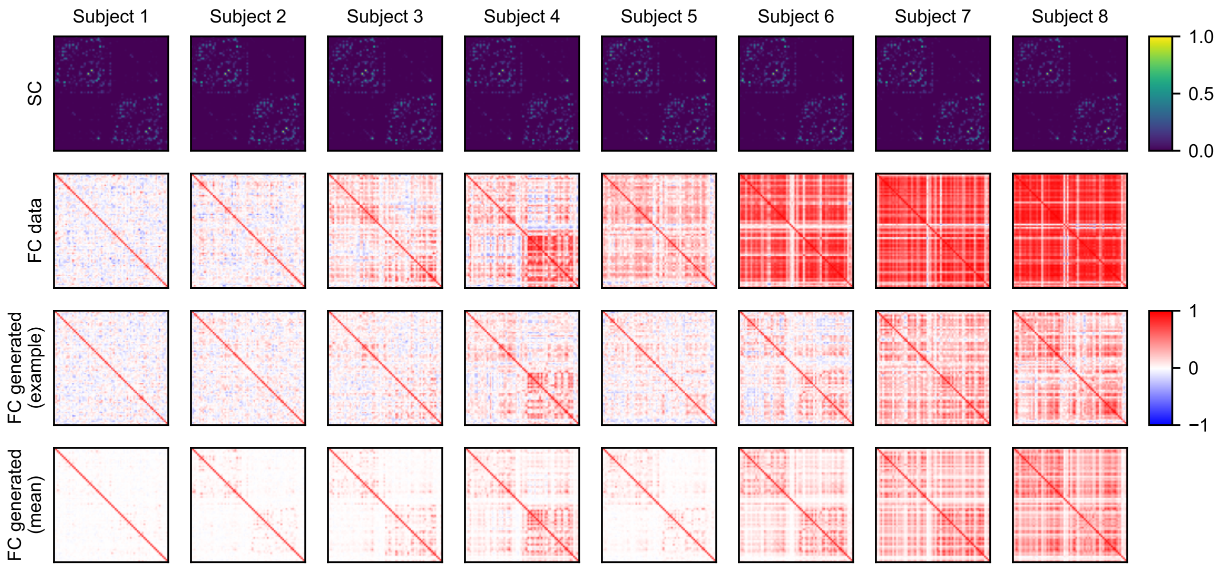


Figure S3: Structural and functional connectivity matrices for all subjects in the Hopf model test case. First row: structural connectivity. Second row: Functional connectivity of the original data used for the training. Third row: Functional connectivity of the example data generated with the trained model. Fourth row: Functional connectivity of the data generated with the trained model, averaged over 50 samples.

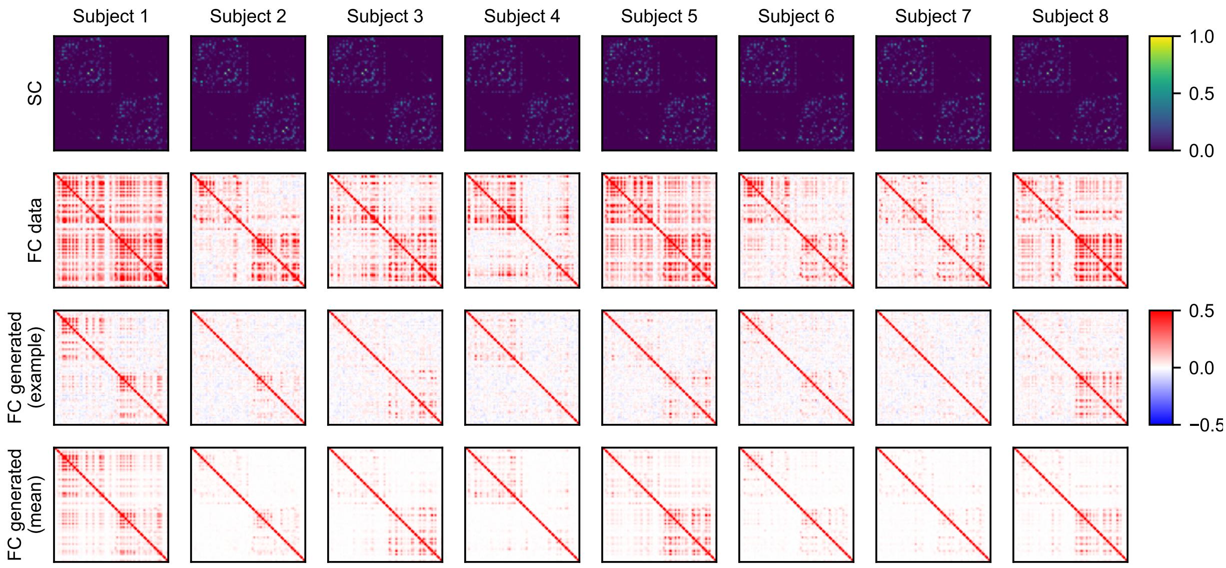


Figure S4: Structural and functional connectivity matrices for all subjects in the pMFM test case. Layout the same as in Fig. S3

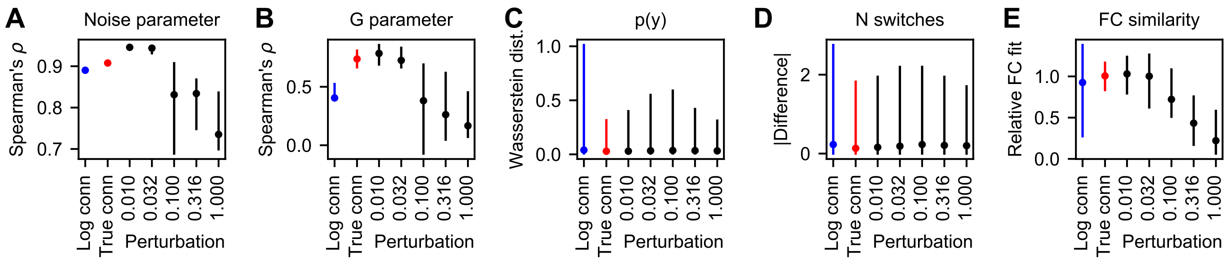


Figure S5: Effect of the connectome perturbation. (A) Spearman's correlation coefficient for the recovery of the noise parameter for the log-scaled connectome (blue), original connectome (red), and five perturbed connectomes (black). (B) Spearman's correlation coefficient for the recovery of the coupling parameter $G$. (C) Wasserstein distance of the distributions in the observation space of the original data and the data generated by the trained model. (D) Difference of the logarithm of number of switches between down- and up-state of the regional timeseries. (E) Relative FC fit, that is, normalized Pearson's correlation coefficient between the non-diagonal elements of the original FC and the FC generated by the trained model. The normalization is performed by dividing the coefficients by the mean of values obtained for the true connectome for every subject separately. The normalization is done in order to make the values comparable across subjects. For all panels, data were generated using four different connectome perturbations for each magnitude value, and one connectome for the original and log-scaled connectome. In panels A and B, 100 samples were drawn from the parameter distributions for each trained model. In panels C-E 50 simulations were performed to calculate the measures of goodness-of-fit for each model. These were then aggregated across all subjects (and across regions apart from panels B and E). Each line represents the 5 to 95 percentile range, with the dot representing the median.

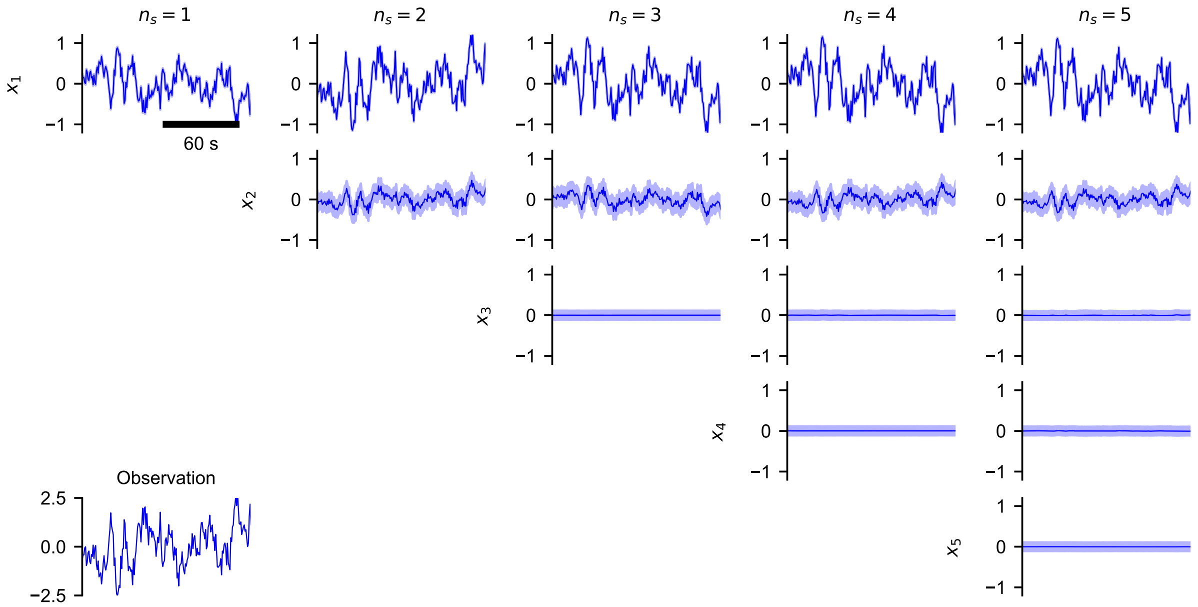


Figure S6: Example of the projection of the observed time series into the state space for models with varying state space dimension $n_s$. In the bottom left corner an observation time series from one region is shown. The columns show the inferred states $x_1$ to $x_5$ when models with state space dimensionality $n_s = 1, \ldots, 5$ are used. For the ease of comparison, the dimensions in each column are order by the amount of information stored in each dimension. The plots show the inferred mean (line) and the span of one inferred standard deviation (shaded area) in each time point. The plots show that in dimensions $x_3$ and higher no information is encoded irrespective of the model state space dimension.

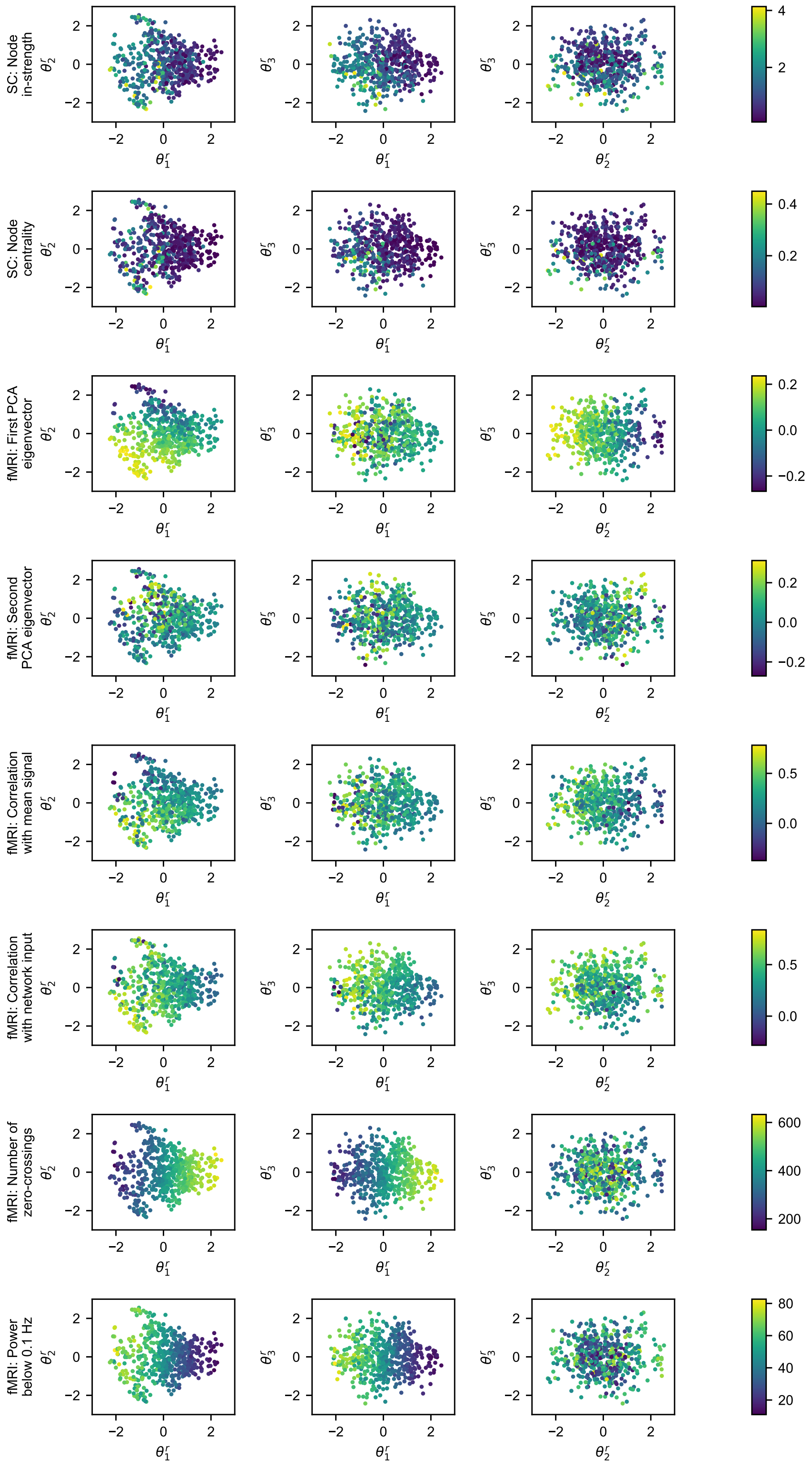


Figure S7: Human resting-state fMRI: Relation between the inferred parameters and features from individual data. Each dot corresponds to one region of one subject. The position corresponds to the inferred mean of regional parameters, the variances are not visualized.

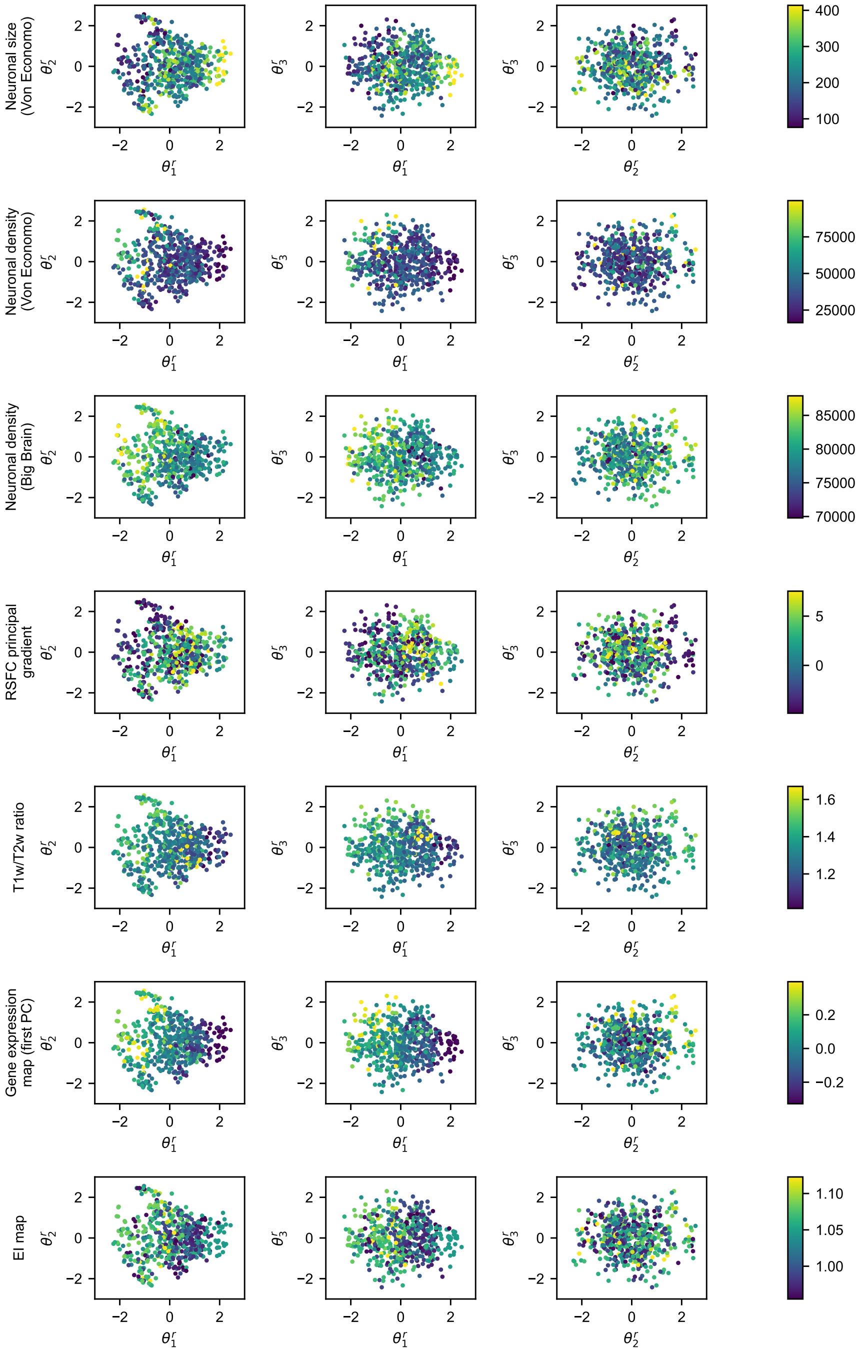


Figure S8: Human resting-state fMRI: Relation between the inferred parameters and features from external data. Each dot corresponds to one region of one subject. The position corresponds to the inferred mean of regional parameters, the variances are not visualized.

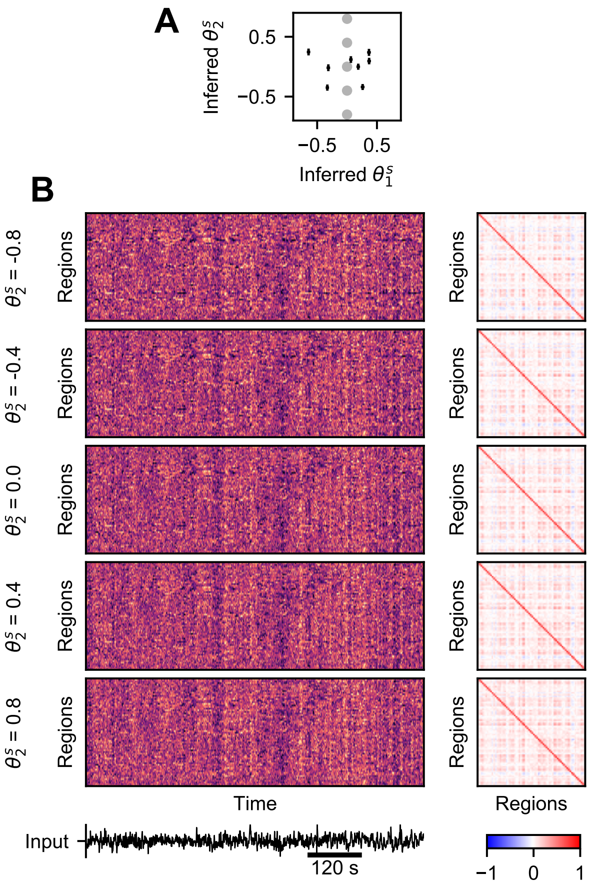


Figure S9: Effect of the subject-specific parameter $\theta_2^s$ on generated whole-brain dynamics. Layout the same as in Fig. 7 in the main text.
